# Mutation distribution density in tumors reconstructs human’s lost diversity

**DOI:** 10.1101/773317

**Authors:** José María Heredia-Genestar, Tomàs Marquès-Bonet, David Juan, Arcadi Navarro

## Abstract

Mutations do not accumulate uniformly across the genome. Human germline and tumor mutation density correlate poorly, and each is associated with different genomic features. Here, we analyze the genome-wide distribution of mutation densities in human and non-human Great Ape (NHGA) germlines as well as human tumors. Strikingly, non-human Great Ape germlines present higher correlation with tumors than the human germline does. This situation is mediated by a different distribution in the human germline of mutations at non-CpG sites, but not of CpG>T transitions. We propose that the impact of ancestral and historical human demographic events on human mutation density leads to this specific disruption in its expected genome-wide distribution. Tumors partially recover this distribution by the accumulation of pre-neoplastic-like somatic mutations. Our results highlight the potential utility of using Great Ape population data, rather than human controls, to establish the expected mutational background of healthy somatic cells.

## Introduction

Mutation density, at different scales, has been shown to correlate with different genomic features, such as regional GC-content or recombination rate^1–5^. In cancer, mutation density has been linked to chromatin states^6^, with higher mutation accumulation in closed chromatin. It has been suggested that the tumor’s higher mutation accumulation in closed chromatin is due to poorer accessibility or recruitment of the mismatch repair machinery to late-replicating, closed-chromatin regions^7,8^. Recent studies have shown that the correlation between tumor mutation density and chromatin state is highly tissue-dependent, allowing the identification of the tissue of origin of metastatic tumor samples^9,10^.

At a smaller scale, sequence context is a good predictor of the mutation rate^11^, beyond hypermutable CpG sites^12–15^. Sequence context has been widely used in cancer analyses to detect signatures of mutation associated with mutagens such as UV-light, tobacco smoke, or APOBEC activity^16,17^. These effects have also been detected in healthy somatic tissues^18,19^.

*De novo* mutations are also affected by sequence context^20–22^. The rates of some particular mutation types have changed recently across ancestries^23–26^. Mutation rates seem to have been under selection in the human lineage. Sequence context studies have shown differences in the relative proportion of certain mutation types between Great Ape species^25^. Furthermore, studies of *de novo* mutations in Great Ape samples revealed a slowdown of the overall mutation rate in humans relative to chimpanzees and gorillas^27^.

Here we study mutation rate evolution, through the differences in mutation distribution (at the 1Mbp scale) between human tumoral tissues and healthy populations in the Great Ape lineage.

## Results

We compared the mutation density distribution in human (1kGP^28^, sgdp_50^29^), non-human Great Apes (NHGA: chimpanzee^30,31^, gorilla^30,32^), and human cancer^33^ datasets. We focused on high-quality orthologous regions shared between human, chimpanzee and gorilla genomes, measuring the number of variants per 1Mbp independently of the frequency of each variant (see Methods).

In agreement with previous reports^1,3,4,6^, we observe a variable distribution of the mutation density across the genome in all datasets (**Figure 1a**). Mutation densities correlate weakly between the human germline and tumors^1,6^ (**Table 1**). Strikingly, the NHGA-tumor correlations are much stronger than the human-tumor correlation and are similar to the human-NHGA germline correlations (**Table 1** & **Supplementary Table 1)**.

**Table 1:** Correlations. Pairwise Pearson’s correlation R of the standardized mutation density of 5,040 1Mbp windows between datasets. Correlation between distributions of mutation density

|  | 1kGP | Chimpanzee | Gorilla |
| --- | --- | --- | --- |
| Chimpanzee | 0.65 | - | - |
| Gorilla | 0.53 | 0.84 | - |
| Tumor | 0.16 | 0.55 | 0.58 |

**Figure 1:**
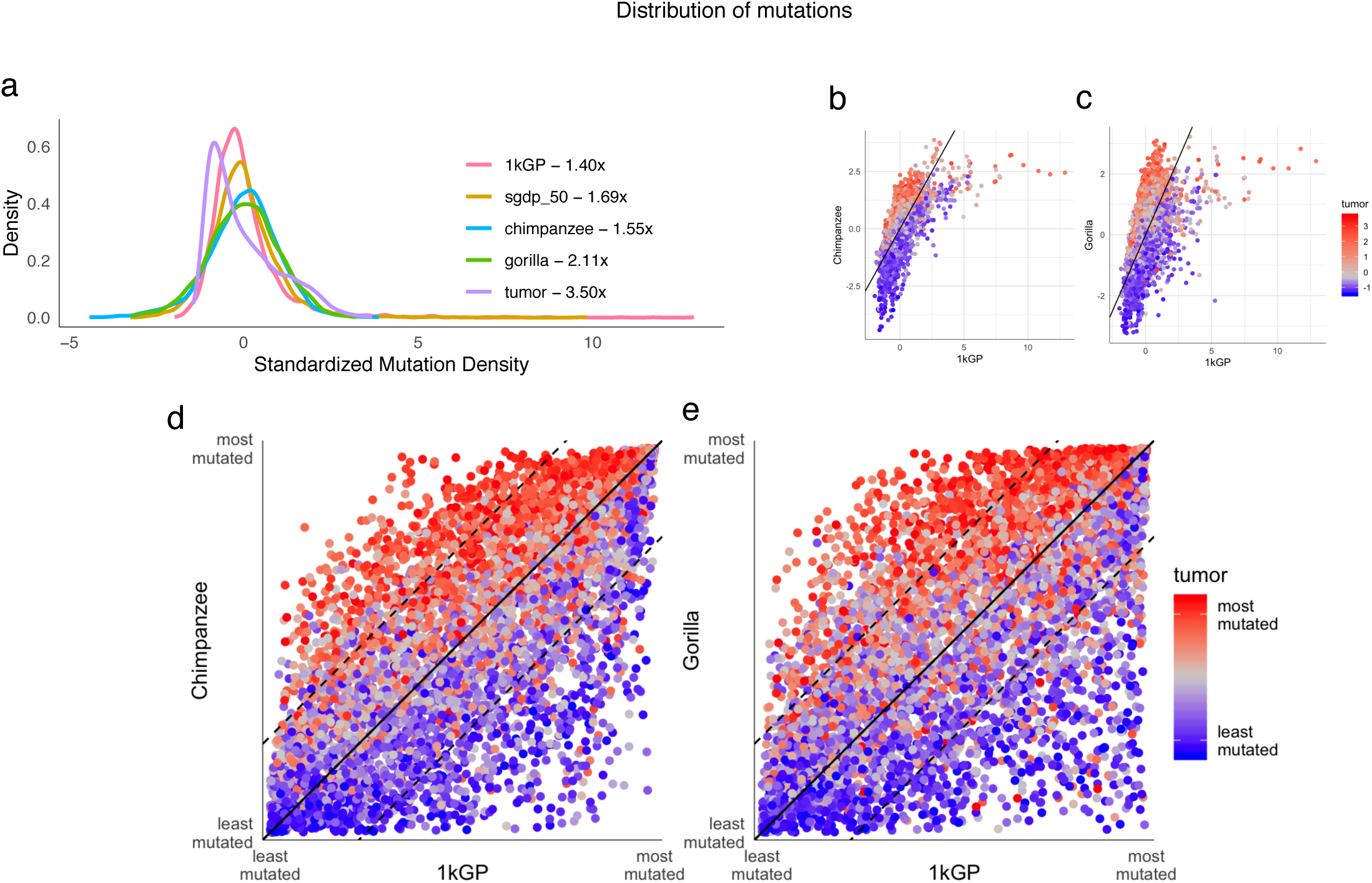
Distribution of mutation density across datasets. **a)** Distribution of the standardized mutation density in 1Mbp windows in human, NHGA, and tumor datasets. The numbers next to the legend represent the fold-enrichment between the 95^th^ and 5^th^ quantiles. **b)** Distribution of the standardized mutation density in humans, chimpanzee and tumor. Each point represents a 1Mbp window. The x-axis represents the human mutation density, the y-axis the chimpanzee mutation density, and the point color, the tumor mutation density. The black line represents the diagonal where the mutation density is equal in human and chimpanzee. **c)** Same as b but comparing human and gorilla. **d)** Distribution of the ranked mutation density in humans, chimpanzee and tumor. Each point represents a 1Mbp window. The x and y axis represent the ranking in mutation density in human and chimpanzee, respectively. Color of points represents the ranked mutation density in the tumor dataset. The solid black line represents the diagonal where the ranked mutation density is equal in human and chimpanzee. The dashed lines represent 25% difference in ranking in both species. **e)** Same as d, comparing human and gorilla.

We compared the distribution of mutation density between pairs of datasets (**Supplementary Figure 1**). Interestingly, we observed that mutation density in tumors is higher in windows where NHGAs have higher mutation density than humans (**Figures 1b,c**). To control for differences in the shapes of distributions, we ranked each set of windows according to their mutation density (**Figures 1d,e**). These ranked distributions show a clear pattern: tumor mutation densities are higher in windows with higher ranks in NHGAs than in human (two-sided Mann-Whitney U test p-value human-chimpanzee= 3.7e-216; human-gorilla = 2.8e-161). This behavior is exclusive to human-NHGA comparisons, as it cannot be observed when comparing chimpanzee to gorilla (**Supplementary Figure 1**), and can be replicated under different conditions and datasets (**Supplementary Notes, Supplementary Tables 2-6 & Supplementary Figure 1**).

High-diversity NHGA subspecies have stronger correlations with both human and tumor than the low-diversity subspecies (**Supplementary Table 3**). Furthermore, the diagonal pattern is only characteristic of comparisons between the germlines of humans and high-diversity NHGA subspecies. A comparison of high and low-diversity chimpanzee and gorilla subspecies showed a clear horizontal split (**Supplementary Figure 2**). Mutation density in tumors co-localizes with the most diverse NHGA subspecies, regardless of the mutation density in the least diverse. In other words; while a lack of diversity distorts the distribution of the genome-wide mutation densities, the diagonal pattern is caused by effects intrinsic to the human lineage. We observed a weak intermediate pattern when comparing NHGA to three archaic hominid genomes (**Supplementary Figure 1**; **Supplementary Note**). This suggests that at least part of the differentiation process in the distribution of mutation densities was already established before the human-Neanderthal split.

Interestingly, correlations between a variety of genomic features and tumor mutation density are consistently more similar to the correlations with NHGAs than with humans (**Figure 2a)**. Mutation densities in NHGAs have, like in humans, strong correlations with sequence conservation and recombination rate (**Supplementary Figure 3**). However, and strikingly, NHGAs show strong positive correlations with epigenomic features associated with closed chromatin, just as tumors do (**Figure 2a, Supplementary Table 7**). We also observe consistent associations with human chromatin states^34^ (**Figures 2b,c**). GC-content, H3K36me1, and CpG-content show a clear positive correlation with human but negative with NHGAs and tumors, suggesting that they might be contributing to the diagonal pattern (**Figure 2d,e** and **3a,b**). Interestingly, H3K36me1 has been shown to be specifically recruited in the gene bodies of genes regulated by CpG islands although its role remains unclear^35^.

Intrigued by the connection of several CpG-related features with the diagonal pattern that implies stronger correlation between mutation densities in tumors and NHGA than with the human germline (**Figure 3a,b**), we analyzed separately CpG>T transitions and mutations at non-CpG sites. (**Figure 3c-f**). CpG>T transitions present very strong correlations between all germline datasets and very poor correlations with tumor (**Figure 3c,d**). The relationship between CpG-content and mutation density at non-CpG sites is different in humans compared to NHGAs and tumors. Moreover, their correlations are similar to those using all sites (**Figure 3e,f**). Correcting the mutation density of CpG>T transitions by the regional CpG content homogenizes the directions of the correlations with genomic features in all datasets (**Supplementary Notes, Supplementary Figure 2**). Interestingly, this correlation is weaker in human than in NHGA and in tumors (**Supplementary Notes, Supplementary Table 8, Supplementary Figure 3**). This suggests that the differences in correlations with genomic features are caused by differences in the relative contribution of non-CpG/CpG>T mutation density in each dataset. The distribution of human *de novo* mutations^36^ at both non-CpG and CpG sites replicates the behavior of human germline mutations showing very low correlations with tumor (**Supplementary Notes, Supplementary Tables 3&9**). When comparing the distribution of non-CpG mutations, we detect a horizontal pattern (**Supplementary Figure 3**) similar to those observed in comparisons of high- and low-diversity subspecies. Therefore, the combination of the behaviors of both non-CpG and CpG>T mutations causes the diagonal pattern observed when comparing all SNVs.

**Figure 2:**
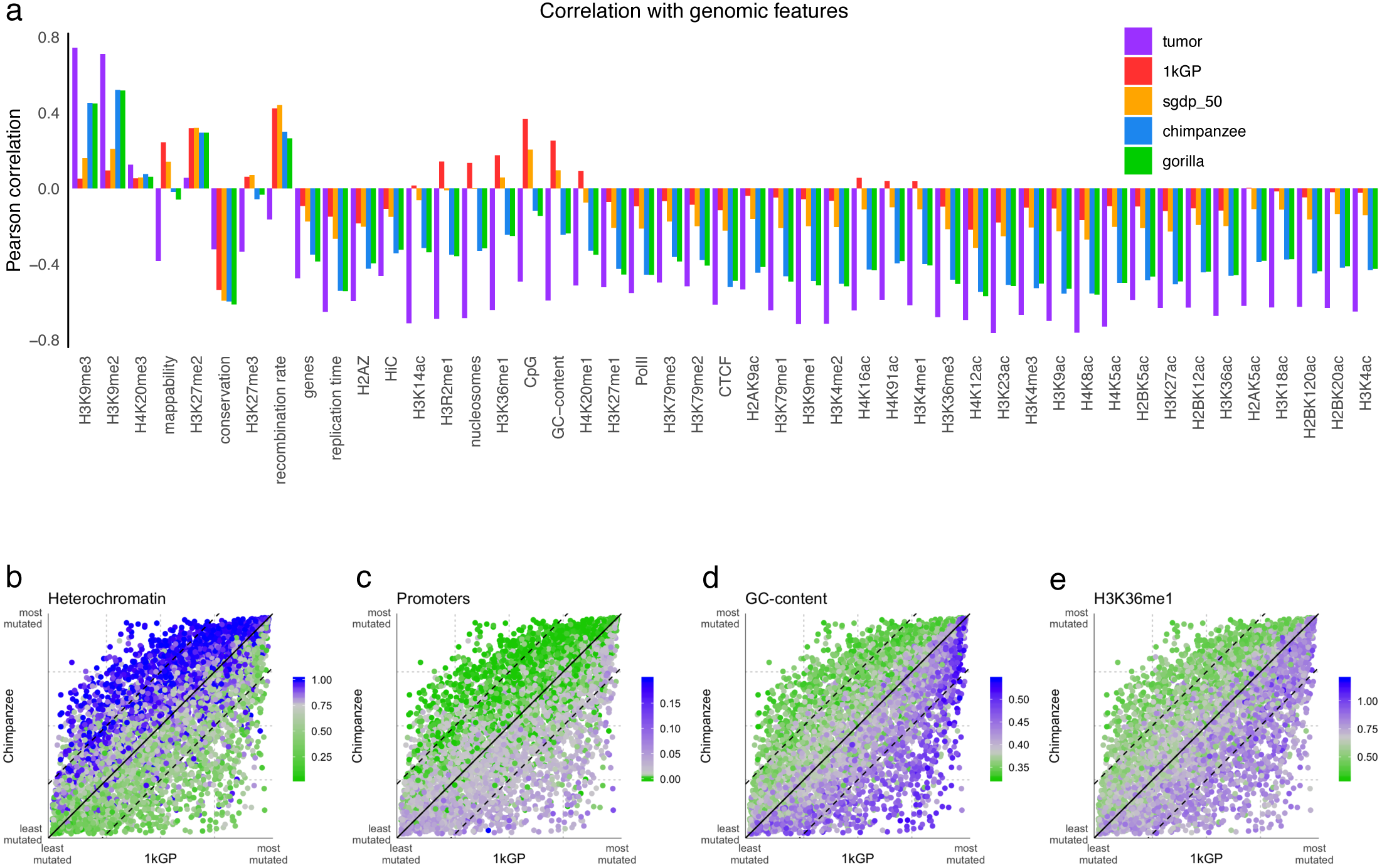
Genomic Features. **a)** Pearson’s correlation R of different datasets with human genomic features (**Supplementary Table 7**). **b)** Overlap of heterochromatin in human lymphoblastoid cell lines (LCLs) measured by chromHMM states^34^ compared with the human and chimpanzee ranked mutation density distribution. **c)** Same as b but using the aggregate chromHMM states associated with the presence of promoters. **d)** same as b and c but color denotes the window’s GC-content, **e)** density of H3K36me1 histone mark ChIP-seq reads^54^.

**Figure 3:**
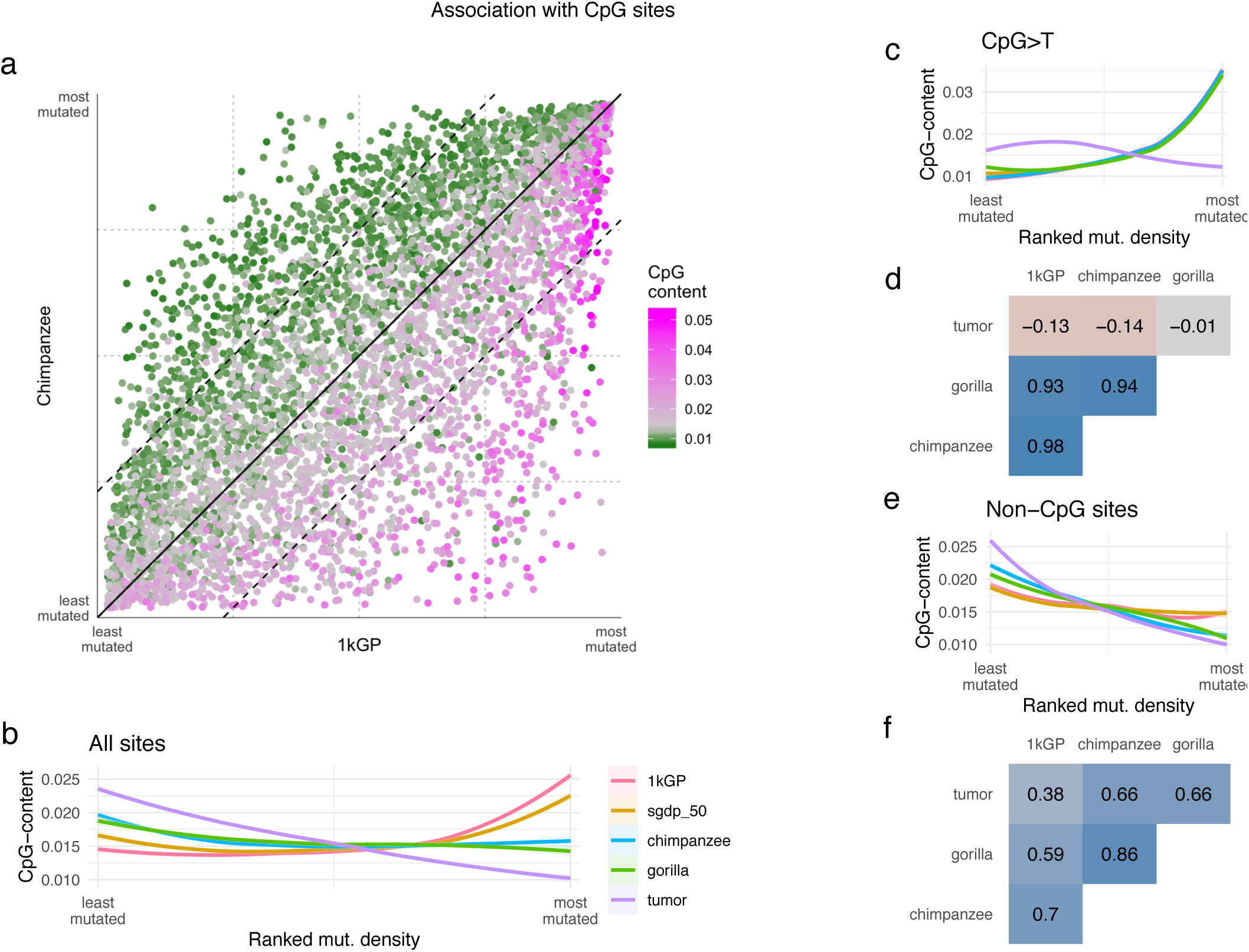
CpG-content**. a)** Distribution of the CpG-content in the human reference hg19 compared with the ranked mutation density in human and chimpanzee, **b)** loess smoothers of mutation density rank and CpG-content for the different datasets. **c)** CpG>T transitions corrected by the whole window size; loess smoothers same as in b; **d)** correlation of the standardized mutation density of CpG>T transitions in different species; **e)** same as in b,c, but using only mutations at non-CpG sites; **f)** correlation of the standardized mutation density of mutations at non-CpG sites in different species.

To explore the contribution of different mechanisms to the observed mutation densities, we analyzed their trinucleotide context. The triplet mutation spectra of human, chimpanzee, and gorilla are very similar (**Supplementary Figure 4, Supplementary Table 10**). It has been shown that the human mutation spectrum can be recapitulated by a combination the cancer signatures SBS1 and SBS5^20,37^. We were able to replicate this association in NHGA and another primate species (Vervet monkey) (**Supplementary Notes, Supplementary Table 10**), suggesting its conservation in the primate lineage.

A subset of trinucleotides is significantly enriched in one of the species (Chi-Squared test p-value <10e-5). We detected no association between these trinucleotides and known mutation mechanisms (**Figure 4a, see Methods, Supplementary Note, Supplementary Figure 4, Supplementary Table 11**). However, linear regression models show a positive and significant (p-value <10e-4) effect of the triplet’s GC-content and its fold-enrichment in the human-chimpanzee comparison (**Supplementary Figure 4**). Only trinucleotides with similar enrichment between species (non-CpG, mainly C>G and T>C) show differences in their distribution across the genome between human, NHGA, and tumor (trinucleotide-difference test p-value <10e-5, see Methods, **Supplementary Note, Figure 4a**).

**Figure 4:**
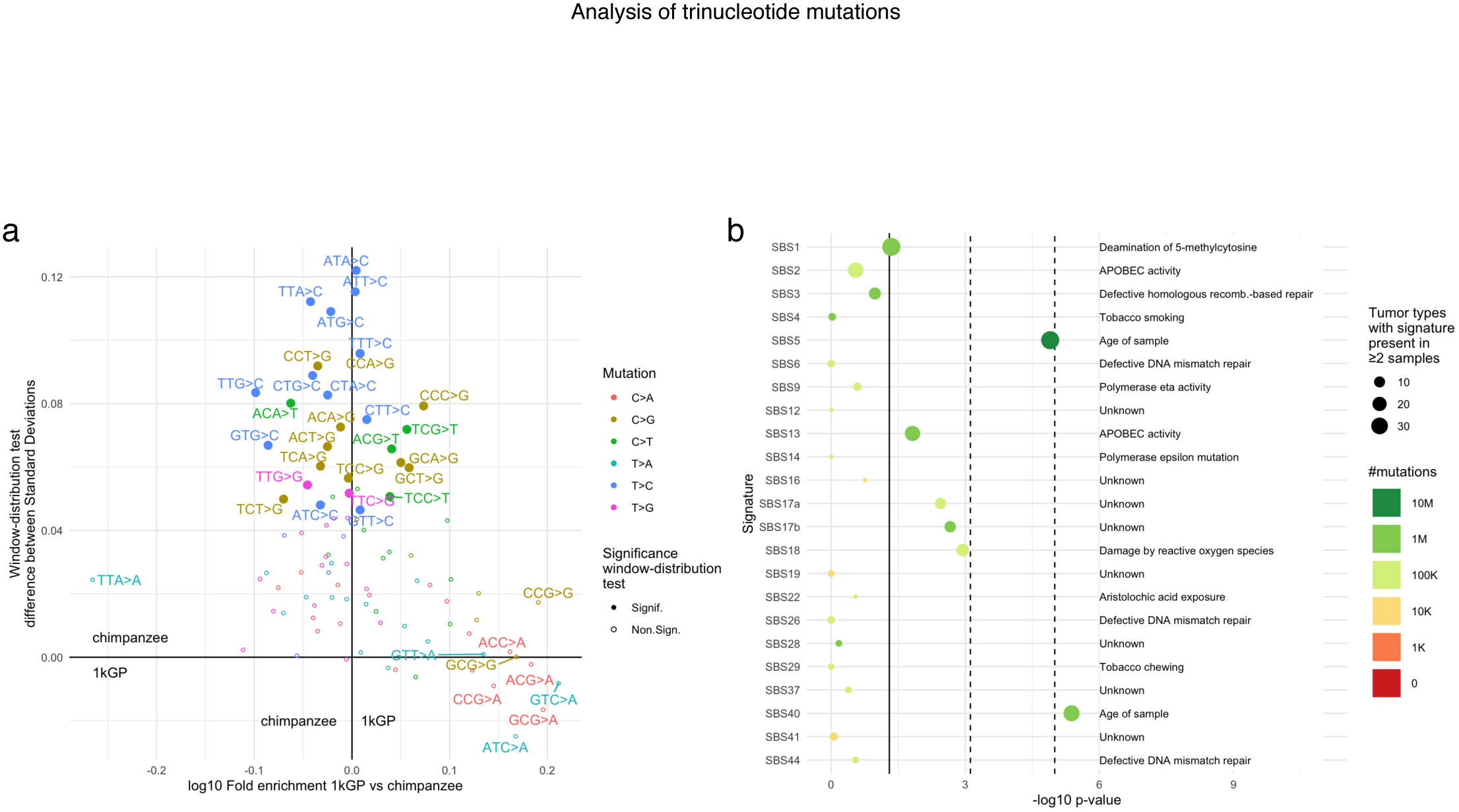
Trinucleotide analysis. **a)** Contribution to the higher chimpanzee-tumors mutation distribution similarity Vs. genome-wide enrichment in human compared to chimpanzee. X-axis: log10 of the enrichment of trinucleotides comparing human and chimpanzee. Left: enriched in chimpanzee; right: enriched in human. Y-axis: effect size (difference between the standard deviations of human-tumor and chimpanzee-tumor) of the trinucleotide-difference test (see Methods). Positive values: tumor distribution more similar to chimpanzee; negative values: tumor distribution more similar to human. Color represents the central nucleotide mutation type. Filled dots represent mutation types significant (p-value <1e-5) in the trinucleotide-difference test. **b)** -Log10 p-values of the association of each cancer signature mutation load to the trinucleotide-difference test (signature-difference test; see Methods). Color represents the number of mutations associated with each signature in the whole dataset. Dot size represents the number of tumor types with two or more samples showing the signature. Only non-artifact signatures present in 2 or more tumor types are shown.

We compared the association of the number of mutations caused by each cancer signature^17,38^ in each individual tumor type to the human-NHGA-tumor pattern (**Supplementary Table 12**). Signatures SBS5 and SBS40 showed a significant association (signature-difference test p-value <10e-4, see Methods) of the pattern with the tumor’s signature mutation load (**Figure 4b**). Both SBS5 and SBS40 are flat signatures whose mutation load is associated with the age of the sample and with pre-neoplastic mutations in tumors^17,38^ This suggests that the strong correlation between NHGA and tumor mutation densities is driven by conserved mechanisms in healthy cells in the Great Ape lineage, while the genome-wide distribution of mutations has been altered in the human germline.

## Discussion

We analyzed the mutation density distribution at the 1Mbp scale in the human and NHGA germlines, as well as in human tumors. We observed a moderate similitude between human and NHGA germlines and, surprisingly, a higher resemblance between human tumors with the germlines of NHGAs than with humans

These discrepancies in mutation density in the human and NHGA germlines are differently associated with genomic and epigenomic features. Regions more densely mutated in humans than in NHGAs tend to be GC-rich, exon-rich, promoter and enhancer-rich, open chromatin and early replicating. Particularly, CpG-related features show a positive correlation with human and a negative correlation with NHGA and tumor mutation densities. The possible functional implications in human evolution require further study.

These observations are driven by the different behavior of mutation density at CpG>T transitions (very similar in all germlines and very different in tumors) and at non-CpG sites (more similar in NHGAs and human tumors than in human germline). This is exclusive of the human germline and, thus, must have been caused by human-specific conditions.

We observed that human and other primates showed a very similar global triplet mutation spectrum. We detected an enrichment of certain trinucleotide mutations in humans and NHGAs consistent with previous results (non-CpG, GC-rich mutations are enriched in humans)^25^. The enriched trinucleotides are not associated with mutation signatures with known causes, nor do they contribute significantly to the higher similitude of human tumors to NHGA germlines. This suggests the absence of strong mechanistic changes biasing the accumulation of mutations in any of the studied germlines.

As previously described for human^20,37^, we observed that mutation rates of three non-human Primates are explained by mutation signatures SBS1 (mostly CpG>T mutations) and SBS5 (associated with “normal” accumulation of mutation in healthy somatic and germline cells^16,39^). Moreover, the lower human-tumor than NHGA-tumor correlation is driven by the accumulation of mutations associated with signatures SBS5 and SBS40 (similar to SBS5 and recently discovered^17^). These results suggest that the poor human-tumor correlation is caused by the fact that human (but not NHGAs) germline (and *de novo* mutations) do not currently reflect the expected mutation densities of healthy (and pre-neoplastic-like) human somatic cells. One possible explanation of this effect, would be if the recent slowdown in mutation rates in humans^27^ affected differently the different types of mutations.

We observed that the moderate human-NHGA and the low human-tumor correlations of mutation densities at non-CpG sites could be caused by losses in population diversity (as observed in low-diversity NHGA subspecies). We propose that successive bottlenecks during human evolution removed a substantial part of nucleotide variation that still remains to be recovered as a whole. In contrast, the hypermutability of CpG sites and its concentration in specific regions caused CpG>T transitions to have already recovered diversity levels similar to those of high-diversity NHGAs. Moreover, the recent human-exclusive population expansions^30,40^ are expected to cause an increase of clock-like CpG>T mutations in the population^41,42^, leaving signatures akin to positive selection, as it has been described in Native Americans^24^. These effects caused a decoupling of the CpG>T/non-CpG mutation rates within the same region, stronger in humans than in NHGA and tumors. We cannot disregard an additional contribution of human-specific shifts in CpG>T transitions mutation rates, although they have been suggested to be similar across all Great Apes^42^. We propose that the combination of population bottlenecks and expansions, together with the specific nature of the different mutation types, drives the differences observed in the distributions of human mutation densities.

Our results imply that accumulated mutations in human populations are a poor proxy of the expected mutational background in healthy somatic cells. In fact, accumulated mutations in NHGAs (at least at non-CpG sites) or even in tumors happen to be more informative about the normal occurrence of mutations in healthy somatic cells.

## Methods

### Datasets used

For the human datasets we used the release variant calling of 2,504 humans from the 1000 Genomes Project^28^ (1kGP), our own calling of 50 additional human samples from the Simons Genome Diversity Panel^29^ (sgdp_50), and *de novo* mutations from 1,548 trios^36^ that were mapped to the human reference hg19 using the liftOver tool^43^. We used our own mapping and calling of 69 chimpanzees and bonobos (59 chimpanzees and 10 bonobos, referred as chimpanzees in short)^30,31^ and 43 gorillas^30,32^. We used the release variant calling of 3 archaic samples: Altai and Vindija 33.19 Neanderthals^44,45^, and Denisova^46^. Finally, for the tumor dataset, we used the release variant calling of 2,583 human tumors from the Pan-Cancer Analysis of Whole Genomes Consortium^33^.

### Definition of high-quality orthologous regions shared between human, chimpanzee and gorilla genomes

We mapped and called chimpanzee and bonobo, gorilla, and human (sgdp_50) samples to the human reference hg19 using BWA MEM^47^ and GATK^48^ following the best practices protocols^49,50^ and additional quality filters (**Supplementary Notes**).

To avoid missmappings to the human reference and erroneous estimates of mutation density in the NHGA samples (too low density caused by lack of mapping reads or deletions or too high density caused by collapsed duplications^51^) we filtered out any region of the human reference genome hg19 failing one of the following criteria: poor mappability of the human reference split into 35bp k-mers, poor callability in ≥25% of the chimpanzee or gorilla samples, or, matching a known Copy Number Variable region in NHGA samples^52^ (**Supplementary Notes**). 2,052Mbp of autosomal sequence passed this filtering (76.54% of the non-N human reference autosomes). We divided the autosomes into 1Mbp overlapping (500kb) windows and kept all windows where ≥50% of its bases passed our filtering. This left 5,040 1Mbp windows to analyze (**Supplementary Figure 1, Supplementary Table 2**).

These filters were applied to all datasets used, including both our callings and external datasets used as released. All SNV counts, trinucleotide counts, and genomic features measurements through this study used only regions passing this filtering. For the analysis of archaic samples, we combined this filtering with the intersection of the callability mask of all 3 archaic samples. This specific filtering was applied to all datasets when compared with the archaic samples.

### Mutation density

We measured mutation density of each window in each dataset by counting either the number of non-fixed segregating sites (in the human, chimpanzee and gorilla datasets) or the number of somatic mutations (in the tumor and human *de novo* datasets, accounting repeated mutations as independent mutational events). We divided this count of Single Nucleotide Variants (SNV) by the fraction of the window passing our filtering. This results in a measure of mutations per Megabasepair (Mbp) of sequence for each window. We standardized the resulting distribution within each dataset deeming it as the mutation density. We ranked all windows within a dataset by their distribution of mutation density to control for the different shapes of the datasets distributions.

### Correlations between distributions

All correlations used in this analysis are Pearson’s correlation (using the R function cor.test) between the standardized mutation densities (unranked) of the two datasets unless otherwise specified. Partial correlations, when used, were calculated using the pcor function from the ppcor R package.

### Significance of the diagonal split

To measure the significance of the diagonal split pattern observed when comparing the human and NHGA datasets, we divided all windows into two groups depending on if the ranked mutation density is higher in human than NHGAs or vice-versa. We calculated the two-sided Mann-Whitney U test on the variable of interest (usually, the tumor mutation density) on both groups using the R function wilcox.test.

### Genomic Features

The genomic features used were filtered using the same mappability, callability and copy-number filters used for the mutation density data. The features used were either the overlap of the feature’s genomic coordinates with the fraction of the 1Mbp window passing our filtering (e.g. GC-content, CpG-content), or the average value or intensity of the feature in the passing fraction of the window (e.g. histone marks), depending on the original data (**Supplementary Table 7**).

### Trinucleotides

We classified each SNV into the 96 possible combinations of trinucleotides (12 different mutation types, by 16 combinations of the adjacent nucleotides, divided by two when folding them). We determined the adjacent reference sequence of each SNV using the getfasta option of bedtools^53^. We filtered out any variant where the liftOver tool^43^ could not map them to the chimpanzee panTro5 or the gorilla gorGor5 reference genomes, or the trinucleotide sequence differed in one of the three reference genomes. This filter was applied to all windows and we used for our analysis only windows where ≥50% of it passed both the original high-quality orthologous regions and this 3-reference filter, leaving 4,920 windows to use. We applied additional filters requiring the trinucleotide to be species-exclusive and to not overlap variants in other species (**Supplementary Note**). This resulted in a high-confidence set of species-exclusive trinucleotides where the ancestral and derived alleles could be reliably inferred. This filtering affected more CpG>T than non-CpG sites, due to the recurrent nature of CpG>T transitions (**Supplementary Table 10**).

### Mutation spectra

We calculated each species’ mutation spectra as the fraction of all trinucleotides in a dataset belonging to one of the 96 trinucleotides. We calculated correlations between datasets using Pearson’s correlation (cor.test function in R). We measured the correlation of the mutation spectrum of each species and the combined effect of cancer mutation signatures SBS1 and SBS5^17,38^ by the formula: 0.1*SBS1+0.9*SBS5, as CpG>T transitions are the main components of signature SBS1 and they represent ∼10% of the trinucleotides in both the human and NHGA datasets.

### Whole-genome enrichment of trinucleotides

We calculated the enrichment and its significance in each germline dataset pair (human-chimpanzee, human-gorilla, chimpanzee-gorilla) using the method described in Harris, 2017^25^. We calculated the enrichment of trinucleotide T between species A and B by dividing *fraction of T in species A / fraction of T in species B*. We calculated a chi-squared test using a contingency table with: the trinucleotide count in species A, in species B, the count of the rest of trinucleotides in species A, and in species B. As the counts of trinucleotides are not independent from each other, we sorted all trinucleotides from most to least significant, and rerun the test by decreasing significance order, while removing the previously used trinucleotides from the count of total trinucleotides.

CpG>T transitions are highly affected by the sample size of the datasets. We ran all the tests using both 1kGP and sgdp_50 as the human dataset. We detected incoherences on the significance and direction of the results in two CpG>T trinucleotides. We report the results using 1kGP where tests using both 1kGP and sgdp_50 are coherent in both significance and direction of the enrichment. The top 10% most enriched trinucleotides in each species pairwise comparison were compared with cancer mutation signatures^38^, and reported when the trinucleotide represented ≥5% of the mutations within a signature.

### Trinucleotide distribution difference test (trinucleotide-difference test)

We developed a method to determine which trinucleotides contribute significantly to the difference between NHGAs-tumors and human-tumors mutation density correlations: For each trinucleotide T and each pair of species (human-chimpanzee, human-gorilla, and, chimpanzee-gorilla) we, subtract the ranked mutation density of T in species A minus the ranking in tumor, and in species B minus tumor. We calculate the two-sided Kolmogorov-Smirnov test (using the R function ks.test) of the two resulting distributions. We use the p-value of the ks-test as the significance of the test and the difference between the standard deviation of both distributions (as both have a mean of 0) as the test’s effect size. The results when using 1kGP or sgdp_50 as the human datasets are concordant in the direction of the association, but we discarded the sgdp_50 results because the smaller number of SNV (and of each trinucleotide type) results in lower power when using sgdp_50.

### Association of GC-content in the trinucleotide sequence

We counted the number of Cytosine and Guanine bases in each trinucleotide and built a linear regression (using the R function glm). The GC-content of the triplet acted as a predictor of the result of the test (the log10 fold-enrichment in the whole-genome enrichment analysis or the difference between the standard deviation of both distributions in the trinucleotide-difference test).

### Mutation load-difference test per mutation signature (signature-difference test)

In order to determine the contribution of each mutation signature to the difference between NHGAs-tumors and human-tumors mutation density correlations, we rerun the trinucleotide-difference test using the 1kGP and chimpanzee datasets, while using the different individual tumor types (**Supplementary Table 12**). For each trinucleotide, tumor type and mutation signature, we built a linear regression (using R’s glm function) where the mutation load of that signature in that tumor type^17^ predicted the effect size in the trinucleotide-difference test for that tumor type (**Supplementary Note**). For each signature, we built a contingency table where all 96 trinucleotides where classified by whether being significant or not (p-value <0.05) in the trinucleotide-difference test, and the significance of the mutation load in the linear regression model. We ran a chi-squared test on that contingency table and obtained its significance.

## Acknowledgements

We thank B. Lehner, D. Weghorn, and I. Lobón for their insights discussing the analyses, and C. Warembourg for her statistical analysis help. J.M.H.G is supported by MDM-2014-0370. T.M.B. is supported by BFU2017-86471-P (MINECO/FEDER, UE), U01 MH106874 grant, Howard Hughes International Early Career, Obra Social “La Caixa” and Secretaria d’Universitats i Recerca and CERCA Programme del Departament d’Economia i Coneixement de la Generalitat de Catalunya (GRC 2017 SGR 880). D.J. is supported by Juan de la Cierva fellowship (FJCI-2016-29558) from MICINN. A.N. is supported by AEI-PGC2018-101927-BI00(FEDER/UE), MINECO-BFU2015-68649-P (MINECO/FEDER, UE), the Spanish National Institute of Bioinfomatics of the Instituto de Salud Carlos III (PT17/0009/0020), and by FEDER (Fondo Europeo de Desarrollo Regional)/FSE (Fondo Social Europeo).

## Author Contributions

J.M.H.G. performed all the analysis. J.M.H.G and D.J wrote the manuscript. T.M.B., D.J. and A.N. conceived and supervised this work. All the authors read and approved the final manuscript.

## Data availability statement

No new data was generated for this study. All the analyses were performed using publicly available data obtained from their original publications, as referenced.

