## Supplementary Text and Figures for "Mutation distribution density in tumors reconstructs human’s lost diversity"

### Supplementary Notes

### Sample selection

We used 59 chimpanzees (10 *Pan troglodytes ellioti*, 19 *Pan troglodytes schweinfurthii*, 18 *Pan troglodytes troglodytes*, and 12 *Pan troglodytes verus*), 10 bonobo (*Pan paniscus*), and 43 gorillas (27 *Gorilla gorilla gorilla*, 1 *Gorilla gorilla diehli*, 7 *Gorilla beringei beringei*, and 8 *Gorilla beringei graueri*) from the Great Ape Genome Project[1,2]. The gorilla sample Serufuli was excluded because the sample uploaded to the Sequence Read Archive was malformed. We also used the whole release dataset of human samples from the 1000 Genomes Project[3], 50 human samples (24 African, 26 non-African) from the Simons Genome Diversity Panel[4] and 2,583 human tumor samples (selected from 2,583 different individuals) from the Pan-Cancer Whole Genome Analysis Project[5].

### Mapping, calling and filtering

#### Mapping

We mapped the human (sgdp_50) and Non-Human Great Ape (NHGA) samples using BWA MEM[6] to the human reference genome hg19. We used the human reference instead of each species own reference to be able to jointly compare Non-Human Great Ape (NHGA) and human samples, and to take advantage of the more complete human gene models and resources available.

When mapped to the human genome, chimpanzee and bonobo samples cover on average 90.93% of the non-N human reference with good mapping quality and coverage. Gorilla samples cover 90.42%. In order to check how this compares with human samples mapping against their own species' genome, we mapped 10 human germline samples from the PCAWG project[5]. Human samples cover 95.40% of the human reference. These human samples are at higher coverage than the Great Ape samples (mean depth of coverage, humans: 38.70x; chimpanzee & bonobo: 25.59x; gorilla: 19.89x). To test if this difference in mapping is due to the different coverage, we downsampled these human samples to the median coverage of chimpanzee samples (23.7x) and they mapped 95.11% of the human reference. This shows that the difference in mapping is not caused differences in coverage but by the intrinsic sequence divergence between different Great Ape species.

The human genome has many repetitive regions. These regions are difficult to map to, even for human samples. We filtered them out to avoid erroneous variant callings from missmapped or collapsed reads (see sections "mappability" and "copy-number"). When these regions are removed, chimpanzee and bonobo samples map to 97.57% of the filtered human reference and gorillas 96.76%, much closer to that of human samples (high-coverage human samples: 98.72%; chimpanzee-level-coverage human samples: 98.36%).

### Variant Calling

We called variants using the GATK-3.7 Haplotype Caller[7] and applied several filters to remove misscalled variants caused by sequencing errors and potential artifacts. The filters applied were adapted from the GATK best practices guidelines [8,9] and forums (https://gatkforums.broadinstitute.org/gatk/discussion/2806/howto-apply-hard-filters-to-a-call-set ).

Variants removed:

- For all variant types:

    - More than 3 variants called within a 10 bp window
    - All samples total depth of coverage < 4*number of samples
    - Individual genotypes with coverage < 4
    - Genotyping quality < 50
    - Quality by depth < 10
- SNV only:
    - Mapping Quality < 40
    - Mapping Quality Rank Sum < -12.5
    - Read Position Rank Sum < -8
    - Fisher Strand Test > 60
    - Strand Odds Ratio > 4
- InDels only:
    - Read Position Rank Sum < -20
    - Inbreeding Coefficient < -0.8
    - Fisher Strand Test > 200
    - Strand Odds Ratio > 10

After that, we used VCFTOOLS[10] to filter out non-biallelic variants (variants with 2 or more alternative alleles) and variants with more than 20% missing genotypes in the samples of a given dataset (chimpanzee+bonobo, or gorilla).

Finally, to minimize the number of variants erroneously called due by missmappings, collapsed duplications or other events, we removed also:

- Variants where heterozygous samples represented more than 80% of all calls.
- SNV within 5bp of an InDel.
- InDels within 5bp of another InDel.
- Regions where the human reference has poor mappability (see section "mappability").

- Regions where > 25% of the samples of a Great Ape species were not deemed callable by GATK CallableLoci (either by too low coverage, too high coverage, or poor mapping quality).
- Copy Number Variable regions in Great Ape lineages (see section "copy-number").

### Mappability

The human reference contains many repeated regions and regions with high homology that could lead to missmapping of sequence reads. In order to minimize the mapping errors caused by these, we used the SNPable regions software (http://lh3lh3.users.sourceforge.net/snpable.shtml) to filter out these non-uniquely mappable regions. To do this, we split the human reference into 35-bp overlapping k-mers, mapped those k-mers back against the same human reference, and kept only those positions of the reference where the majority of overlapping reads mapped uniquely and without mismatches. This resulted in a list of uniquely mappable regions that cover 80.22% of the non-N human reference genome.

### Copy-Number

In order to avoid erroneous variant calls from collapsed duplications, we used previously published Copy-Number estimates from Great Ape samples[11]. We used the LiftOver tool[12] to convert the coordinates to the hg19 reference. We filtered out a total of 134Mb of sequence in regions defined as fixed duplications in human, chimpanzee, bonobo or gorilla samples, as well as those regions copy-number-variable in at least two samples in any of these species.

### Variant Calling Results

Variant calling resulted in 52.8M SNVs and 1.6M InDels called in the chimpanzee+bonobo dataset (22.2M SNVs and 0.8M InDels per sample on average), and 41M SNVs and 1.6M InDels called for gorilla samples (30.5M SNVs and 1.3M InDels per sample on average). These samples are NHGA mapped to the human reference, and these numbers include all positions fixed in NHGA but different from the human reference (and thus, derived in either the human or the NHGA branches). If we take into account only non-fixed variants, the average number of segregating sites per sample is 5.9M SNVs and 245k InDels in the chimpanzee+bonobo dataset, and 5.8M SNVs and 271k inDels in the gorilla dataset.

When analyzing each NHGA subspecies individually, we observe that NHGA samples have an average number of SNV much higher than human samples. This is caused by between-subspecies differences, as the average number of heterozygous sites per sample of each subspecies is much closer to the human average. This number of heterozygous sites per sample can be used as an estimator of population diversity. The results of our calling show chimpanzee between-subspecies estimates similar to those reported in the original manuscript[13] where these samples were mapped to the chimpanzee reference (Average count of heterozygous SNV per sample: *Pan paniscus*: 1,182,432; *Pan troglodytes ellioti*: 2,013,762; *Pan troglodytes schweinfurthii*: 2,477,963; *Pan troglodytes troglodytes*: 2,999,398; *Pan troglodytes verus*: 1,294,985; *Gorilla beringei beringei*: 1,255,272; *Gorilla beringei graueri*: 1,294,715; *Gorilla gorilla gorilla*: 2,984,205; *Homo sapiens*: 1,669,533).

#### Calculation of mutation density

We divided the autosomes into 1Mbp overlapping windows (with an overlap of 50% or 500kb). We counted the number of SNVs in each windows (segregating sites in the human, chimpanzee and gorilla population -or germline- datasets; and somatic mutations in the tumor dataset). Mutations present in more than one sample were considered as a single mutation event in the germline datasets, but independent mutation events in each sample in the tumor somatic dataset. We divided this resulting number by the fraction of the window passing our filtering, giving a measure of mutations per 1Mbp. Removing recurrent events from the tumor dataset had no effect in the results of the analyses.

### Robustness of the results

#### Effect of the filtering in the windows

The objective of the filtering we used (mappability + callability + copy number) is to analyze only regions of the genome where human and NHGAs have a similar ability to map and call variants. This allowed us to reduce the bias in mutation density estimates in different species. Regions with low homology between humans and NHGAs would show extremely low mutation density in chimpanzee and gorilla because we would not be able to call variants there. The mappability filter has the biggest effect in the global filtering, where 74.07% of the whole genome (including N sites) passes it. The CNV filter removes an additional 1.55% of the genome, and the callability filter of >75% of the chimpanzee+bonobo, and gorilla samples, removes an additional 2.69% of the genome, leaving a total of 69.83% of the genome passing all filters.

Not all the genome is affected equally by this filtering. When dividing the autosomes into 1Mbp windows, we observe a bimodal distribution: some windows pass 0% of our filtering (mainly centromeres, telomeres, and the regions adjacent to them), while most windows pass >50% of our filtering (most passing 75%~90%) (**Supplementary Figure 1**). We set an arbitrary limit of 50% for the windows to analyze, leaving 5,040 windows out of 5,776 for our analysis. The differential distribution of mutation density in human, NHGAs, and tumor is preserved when using higher values for the filtering threshold (**Supplementary Table 2 & Supplementary Figure 1**). When using a filtering threshold of 50%, the human datasets have almost no correlation with the fraction of the window passing our filters (window ratio), while the Great Ape samples show a positive correlation (as expected because one component of this filtering is calculated from these samples). Surprisingly, the tumor dataset shows a strong negative correlation with the window ratio that might be mediated by GC-content or CpG-content (**Figure 2d & 3a, Supplementary Figure 3, Supplementary Table 3**).

When analyzing the effect of the different components of this filtering in the distribution of mutation density, mappability has a high effect across all windows, with slightly more effect in windows with low mutation density in all species or with higher mutation density in humans than NHGAs. Callability has a lower effect than the mappability filtering. We would expect the callability filtering to affect more windows with low mutation density in NHGAs (as we would not be able to call variants in them) while being randomly distributed in human (as they should not affect the human genome). But the results show that NHGAs callability has a strong effect only in the windows more densely mutated in human, independently of the density in NHGAs. This suggests these windows have been somehow recently innovative in humans, and these differences cause the mapping problems in NHGA samples. The CNV filtering does not affect differently windows more highly mutated in one species than the other (data not shown).

We compared the mutation density in both the fraction of each window passing and failing the different components of the filtering. We observe that NHGAs have much lower mutation density in the fraction of the windows failing their own species callability filtering, than in the fraction of the window passing all filters. Meanwhile, in both humans and tumors, the mutation density is similar in both the passing fraction and the fraction failing callability in any NHGA species. The CNV filtering has almost no effect in humans, while filtering out in NHGAs some regions that have either higher or lower mutation density, putatively duplications and deletions. These results show that these filters applied are necessary to remove problematic regions inside the windows that could alter the average mutation density of the window.

### Window Size

We analyzed the distribution of mutation density at different scales using 10Mbp, 100kbp, and 10kbp windows. The correlations between all germline datasets are stable across all window sizes, becoming weaker in 10kbp windows. Tumor has lower correlations with human than with NHGAs across all window sizes (**Supplementary Table 2**). The human-NHGA-tumor pattern can clearly be detected in 10Mbp and 100kbp windows. Using 10kbp windows, this pattern becomes less apparent, but we can observe a clear distribution driven by the higher correlation between NHGAs and tumor (**Supplementary Figure 1**). The Mann-Whitney U tests of the mutation density in tumor are always significant, either splitting the dataset above/below the diagonal, or by windows with >25% difference in human and NHGAs.

### Sample Size

We tested the robustness of the results using subsamplings of the human dataset 1kGP, with the same sample size as the Chimpanzee (69 samples) and Gorilla datasets (43 samples). The samples were chosen aiming to maximize human diversity. We clearly observe similar correlations between the human, NHGAs and tumor datasets (**Supplementary Table 3**), as well as the same pattern in the distribution of mutations where windows with higher mutation density in NHGAs than in humans, have also higher mutation density in human tumors (**Supplementary Figure 1**). This suggests that the different sample size of the datasets is not the cause of the observed association.

### Additional human datasets

We tested the robustness of these results using three additional human datasets different than the 1000 Genomes Project. We used the release dataset of the

International Cancer Genome Consortium (ICGC) germline dataset[5] (the germline samples of some of the tumor samples used in this study), with 1,823 samples; the Simons Genome Diversity Panel (SGDP) release dataset[4], with 279 samples; and our own mapping and calling of 50 samples of SGDP (SGDP_50) chosen to maximize human diversity. We can clearly observe that using these three human datasets shows similar between-species correlations (**Supplementary Table 3**), and distribution (**Supplementary Figure 1**) than when using the 1kGP dataset.

These three datasets, as well as the tumor dataset, were generated using PCR-free technology, in contrast with the 1kGP and NHGAs datasets. Other studies[14] have shown that PCR-amplification causes a decrease in coverage of GC-poor/AT-rich regions in Illumina reads, and this effect should disappear when using PCR-free technologies. We detect differences in the distribution of coverage between the 1kGP and NHGAs datasets, and the PCR-free tumor, ICGC, and SGDP_50 datasets. Having seen that the distribution of mutation density is very similar when using both 1kGP and these three additional human datasets, proves that this differential distribution is robust to different human datasets and it is not dependent on the underlying sequencing technology used.

### Chimpanzee reference

To assess the effect of calculating the distribution of mutation density in NHGAs using data mapped to a reference genome different than their own species, we mapped 50 chimpanzee samples (10 bonobos, and 10 samples from each chimpanzee subspecies), and 50 human samples (SGDP_50) to the chimpanzee reference PanTro5. We ran the same mapping and variant calling pipeline used for the human reference calling. We applied a similar filtering, calculating the unique mappability of 35-mers against the PanTro5 reference, the callability of ≥75% of the chimpanzee and human samples, and the Great Apes CNVs using the LiftOver tool[12] to map them against the chimpanzee reference. We used the number of non-fixed segregating sites in both species to calculate the mutation density in each window. We used LiftOver also to map the tumor dataset to the PanTro5 reference, and calculated the GC-content of each window in the PanTro5 reference (**Supplementary Table 2, Supplementary Figure 1**). We observe the exact same distribution in tumor mutation density and GC-content as in the datasets mapped to the human reference. Windows where chimpanzee has higher mutation density than human, have also higher mutation density in tumors and lower GC-content than the regions with higher mutation density in humans than in chimpanzee. The fraction of the window passing our filtering is no longer associated with low callability in chimpanzee in GC-rich regions, as in the human reference. This proves that this distribution is neither dependent on the reference used to map the samples, nor the fraction of the windows passing the filters.

### Variant Frequency

We divided our datasets and calculated the distribution of the mutation density of each window using only variants of a certain frequency and compared it with the mutation density of the tumor dataset (without altering the tumor dataset variant frequency) (**Supplementary Table 5**). We can clearly detect the differential correlations and distribution pattern in both chimpanzee and gorilla when calculating the mutation density using only non-singleton variants, variants with allele frequency > 5%, variants with allele frequency ≤ 10% or variants with allele frequency > 10%. Variants with allele frequency ≤ 5% do show a clear differential pattern when comparing human with chimpanzee, but the pattern is much more diffuse in gorilla, driven by a lower correlation between gorilla and tumor (data not shown).

When using only singletons or doubletons, the distributions become much more diffuse. The differential correlations between tumor and human or NHGAs are still present. However, the correlations of low-frequency variants between human and Great Ape datasets become much weaker, removing the original human-Great Ape distribution pattern, and, thus, removing the clear diagonal split of the data.

We also removed any shared variant between the human and chimpanzee or gorilla, calculating the mutation density using only variants exclusive of a given dataset (**Supplementary Table 5**). Using only exclusive variants shows clearly the differential pattern, proving that it is not dominated by variants shared between species, but variants that arose in only one of them. The correlation between shared and species-exclusive variants is also similar in the different species (**Supplementary Table 5**).

Previous studies in this field have used human-chimpanzee and human macaque divergence[15–17]. We reconstructed the human-chimpanzee and human-gorilla divergence from population data. We selected all those variants called homozygous for the alternative allele in all samples of a NHGA dataset, that were not found to be segregating in the 1kGP human dataset. We calculated the mutation density of all windows using only these variants (**Supplementary Table 5**). This distribution does not show any clear pattern. The correlation of these variants with all segregating sites is similar in both human and NHGAs. We cannot observe the differential pattern using divergence data. This distribution is heavily affected by the amount of Great Ape samples used, as new low-frequency variants will only decrease the number of fixed sites (increasing from 50 to 69 chimpanzee samples leaves only half of the fixed sites; data not shown). We can extract no conclusion of this, but it would be worth revisiting once more Great Ape samples are available and inferences of divergence from population data are more reliable.

This pattern being observable by variants of high frequency, but becoming more diffuse at extremely low frequencies suggests that there is a common origin in the molecular mechanism or selective pressures generating it, before the human-chimpanzee-gorilla split. This effect can clearly be detected using variants exclusive from the human and Great Ape braches, showing that the mechanisms generating it have been acting individually in each branch after the species split. These mechanisms have been common between human and chimpanzee until very recently, but diverged earlier in gorilla, shown by the lower effect in the accumulation of new low-frequency variants.

Recent studies[18] have detected differences in the mutation rate in human in extremely rare variants and *de novo* mutations when compared with variants at higher frequencies in the population. This suggests that the mechanisms leading to mutation rates have evolved recently in the human genome. NHGA populations could be under similar (although different) effects, leading to this higher differentiation in low-frequency variants.

### InDels

Previous studies[17] have described a co-localization of insertions and deletions in regions with high substitution rate and high GC-content. We analyzed the distribution of non-fixed InDels segregating in the population of each of our datasets (**Supplementary Table 6**). The density of InDels correlates best with the mutation density of its own species. InDels in human and chimpanzee have a poor correlation with mutation density in tumor, while InDels in gorilla show a moderate correlation with tumor. When comparing the InDel densities between species, all germline datasets have moderate pairwise correlations. Tumor InDels have a weak correlation with human InDels, slightly stronger with chimpanzee, and a moderate correlation with gorilla InDels. Comparing the distribution of human, Great Ape, and tumor InDels shows a diagonal pattern only in the case of human-gorilla. This pattern also appears, albeit weakly, when comparing InDel density chimpanzee, gorilla and tumor. This suggests that this co-localization of InDels in gorilla and tumor is either specific from the gorilla lineage or changed towards the current human distribution in the branch of the common ancestor between human and chimpanzee. Another possible explanation could be that the higher divergence time between human and gorilla makes some regions prone to erroneously call InDels instead of more complex structural events. If these regions were to be more prone to a certain degree of structural variation in the human-chimpanzee branch, they could imply also a tendency towards InDel mutations in tumors.

### Tumor types

To determine if this differential mutation density pattern we detect is caused by specific tumor types, we divided 2,581 tumor samples with known histology into 37 histological categories (as reported in the original dataset)[5] and calculated the window mutation density on each of them. We observe clear differences in the correlation of human and NHGAs with each of the tumor types (**Supplementary Table 4**). Although there is variability between different tumor types, with some tumors having a stronger correlation with the germline human mutation density than others, these human-NHGA-tumor differences in correlations are present in virtually all tumor types with enough sample size and number of SNV.

### Archaic hominids

We downloaded the variant calling files from 3 high quality archaic hominids: Altai and Vindija 33.19 Neanderthals[19,20], and Denisova[21]. We downloaded the callability mask of all 3 samples (from http://cdna.eva.mpg.de/neandertal/Vindija/), and intersected it with our previous filters. This left a total of 1,433 Mbp of high quality sequence to analyze, and 4,130 1Mbp windows where ≥50% pass the filtering. Using this new common filtering, we calculated the mutation density in 1Mbp windows in a dataset composed by all 3 archaic samples together, as well as our previous human, chimpanzee, gorilla, and tumor datasets (**Supplementary Table 2, Supplementary Figure 1**). We observe an intermediate correlation between modern humans and archaic hominids mutation density, similar to the modern humans-chimpanzee correlation. This might be caused by real differences in mutation density, or by the extremely small sample size (3 samples) of the archaic hominids. In contrast, the correlation between archaic hominids and tumor mutation density is very low, similar to the human-tumor correlations, and much smaller than the tumor-NHGA correlations. When comparing the ranked mutation density of archaic, we observe a random distribution of tumor mutation rates when comparing modern humans and archaic hominids. Comparing NHGAs with archaic hominids shows a pattern more similar to the human-NHGA-tumor distribution. This pattern does not reproduce the diagonal pattern previously seen and is dominated by the higher tumor-NHGAs correlation, probably caused by the small sample size of archaic hominids. This suggests that the mechanisms causing the differences between humans and NHGAs have been active since well before the split between humans and Neanderthals and Denisovans. We found no association with regions of Neanderthal introgression in modern humans (data not shown).

### Great Ape subspecies

We calculated the mutation density of each NHGA subspecies separately, to assess if specific populations drive this human-NHGA-tumor pattern observed. We observe that, the less diverse subspecies *Pan paniscus* (bonobos), *Pan troglodytes verus* (western chimpanzees), *Gorilla beringei beringei* (mountain gorillas), and *Gorilla beringei graueri* (eastern lowland gorillas), have lower correlations (**Supplementary Table 3**) with human mutation density than the most diverse subspecies (*Pan troglodytes troglodytes* (central chimpanzees), *Pan troglodytes schweinfurthii* (eastern chimpanzees), *Gorilla gorilla gorilla* (western lowland gorillas)). This lower correlation is not mediated by their lower sample size, as subsets of 5 *Pan troglodytes troglodytes* or 5 *Gorilla gorilla gorilla* samples have higher correlations. These low-diversity subspecies also have lower correlations with the mutation density in tumors than the more diverse subspecies, but the correlation between Great Ape subspecies and tumor is always higher than between different human datasets and tumor. Low-diversity subspecies also have lower correlation with GC-content and the fraction of the window passing all our filters.

We compared each of these subspecies against a subset of humans of the same sample size. We selected different subsets of human samples of African origin to maximize human diversity, from the previously used subsets of the 1kGP dataset. We can clearly reconstruct the human-NHGA-tumor pattern when using high-diversity subspecies, even when using only 5 samples in the NHGA dataset. Low-diversity subspecies, though, have a much more random distribution of the mutation rate in tumor, with no clear diagonal pattern, although they have a slightly higher correlation with tumor than humans (**Supplementary Figure 2**). We hypothesize that this diagonal-split distribution is caused by comparing a high-diversity dataset against a low-diversity dataset, regardless of the low-diversity dataset being human samples or not. If this were to be true, we should be able to reconstruct the diagonal split by comparing high-diversity NHGAs with low-diversity NHGAs. When we do this, we observe a clear horizontal split distribution of the mutation rate of tumor, mediated by the higher correlation of tumor with the high-diverse datasets, but we were not able to reconstruct the diagonal split that we see when comparing high-diversity NHGAs with humans. Comparing two low-diversity subspecies shows no clear pattern, while comparing two high-diversity subspecies shows tumor mutation density being higher in the windows with higher mutation density in both subspecies. It is worth noting that, in none of the tested cases, we were able to invert the human-NHGA-tumor pattern and detect higher mutation density in tumors in windows with higher mutation density in humans than in NHGA, even when comparing humans with NHGA subspecies with lower diversity than humans.

### Genomic Features

#### Recombination

Recombination rate has previously been correlated with high mutation density [16,22–24] but has shown to have poor correlation with mutations in tumor[25]. We compared male, female, and average human recombination maps[26], human hotspots of Double Strand Breaks (DSB) in PRDM9 binding sites[27], chimpanzee (*Pan troglodytes verus*) recombination maps[28], and chimpanzee (*Pan troglodytes ellioti*), bonobo and gorilla (*Gorilla gorilla gorilla*) recombination maps[29] (**Supplementary Table 7, Supplementary Figure 3**). All measurements of high recombination rate tend to locate in windows with high mutation density in both humans and NHGA, with no big differences between species.

#### Chromatin State

Previous studies have analyzed the effect of a wide variety of genomic features in the variation of mutation density in the human germline and in human tumors [25]. Conservation and recombination rate were described as the two main drivers of variation in distribution in the human germline, while a wide range of histone marks affecting the chromatin state had very weak correlations with germline mutation density distribution. In contrast, tumor mutation density showed very strong positive correlations with histone modifications associated with closed chromatin, and very strong negative correlations with modifications associated with open chromatin, suggesting that mutation rates in tumor are higher in regions of closed chromatin, probably driven by a poor accessibility to the mismatch repair machinery. Further studies have shown that this correlation between chromatin state and tumor mutation density is more strongly correlated when matching the cell type of both the tumor and the histone marks used[30]. This strong correlation between the cell-of-origin chromatin structure and mutation distribution allows even to distinguish the tissue of origin of metastatic tumors[31].

We reproduced the analysis of Schuster-Böckler, B., Nature 2012 using our human, tumor and NHGA datasets. We used the same list of histone marks, chromatin modifiers, and genomic features from the original article using the same data sources and processing (**Supplementary Table 7**), except an updated measurement of conservation using 100 vertebrates[32] instead of 29 mammals and updated versions of the ENSEMBL gene lists. The results using our datasets match perfectly the results for 1Mbp windows of Figure 1 and Supplementary Figure 2 from Schuster-Böckler, B., Nature 2012 (**Figure 2**). The correlations of both tumor and human germline mutation densities with genomic features show the same trends as the original article. Tumor has a strong positive correlation with closed chromatin features and a strong negative correlation with open chromatin features, while human germline has a strong positive correlation with recombination rate and H3K27me2 (unexplained in the original article), a strong negative correlation with conservation, and lower correlations with the rest of features. The correlations with mappability of our dataset are biased by our previous mappability filtering of all the datasets. We also analyzed the mutation density of human-chimpanzee and human-gorilla divergence measured as fixed SNV called in the Great Ape datasets and not present in the human population. This measurement shows differences with the original article human-chimpanzee divergence, but all the correlations are very weak and in the same direction as the human germline (data not shown). Finally we analyzed the correlation with genomic features of chimpanzee and gorilla mutation densities. Both chimpanzee and gorilla, like human germline, have a strong positive correlation with recombination rate and H3K27me2, and a strong negative correlation with conservation. Strikingly, both chimpanzee and gorilla have moderate to strong correlations with the rest of histone modifications, in the same direction as tumors, although weaker.

We downloaded ChromHMM chromatin state partitioning data from ENCODE[33] in lymphoblastoid cell-lines and compared the overlap of the different chromatin states with the human-NHGA distribution of mutation density. We observe a very similar pattern as previously described, where regions with higher mutation density in NHGAs than in humans co-localize with closed-chromatin features (**Figure 2**).

# GC

We calculated GC-content as the fraction of the base pairs in a window passing our filters and deemed to be either Guanines or Cytosines in the human reference Hg19. We observe a clear differential distribution with GC-content being higher in windows whit higher mutation density in human than in NHGAs (**Figure 2d**), This distribution is consistent when mapping the samples against the chimpanzee reference panTro5 (**Supplementary Figure 1**). It has been previously reported[15,16] that human mutation density (calculated as human-chimpanzee or human-macaque divergence) has a non-linear association with the GC-content of the underlying sequence at 1Mbp scale. We do detect this U-shaped distribution, but it is driven by a very small number of windows with extremely high GC-content and CpG-content. When we plot this inverting the axes and using the ranked mutation density (which has a similar effect to binning windows by their mutation density), humans show a clear tendency of increasing GC-content in high mutation density windows, while NHGA have a slightly decreasing tendency, and tumor a strong decreasing tendency (**Supplementary Figure 3**). The different NHGA subspecies show similar tendencies with GC-content. When comparing the mutation density of high diversity and low-diversity subspecies, we do not observe a clear pattern in the distribution of GC-content as in the case of comparisons with human (**Supplementary Figure 3**). The association of GC-content and the ranked mutation density of NHGA subspecies is not different from that of using all subspecies pooled together. This association also holds when using human and chimpanzee samples mapped to the chimpanzee reference panTro5 (**Supplementary Figure 1&3**). GC-content also correlates with the density of protein-coding exons[34–36] (**Supplementary Figure 3**).

### CpG sites

CpG hypermutable sites have a mutation rate one order of magnitude higher than other sites in the genome[37–40]. CpG-sites correlate highly with GC-content and genomic and epigenomic features.

We filtered our datasets, selecting only those sites where the reference in human (hg19), chimpanzee (panTro5) and gorilla (gorGor3) has the exact same nucleotide in that site and the adjacent +/-1 base pairs. This allows us to compare mutations at CpG sites without biases caused by CpG>T changes accumulated in the branch of the reference used and fixed in their genome. CpG sites show the same differential distribution as GC-content (**Figure 2ab, Supplementary Figure 3**), and represent a similar fraction of all SNV in all germline datasets (**Supplementary Table 8**). Tumor has a lower fraction of mutations at CpG sites.

### CpG methylation

The methylation status of CpGs has an important effect on their mutability, as methylated CpG sites can undergo spontaneous Cytosine deamination, mutating to a Thymine and leading to a transition. CpG-islands are deemed to have lower mutation density, as they tend to be unmethylated. To analyze if the different association of GC-content with mutation density in humans and NHGAs, we downloaded information of the methylation status of human CpG sites from the ENCODE project[41], and differences in methylation of CpG sites in human, chimpanzee, and gorilla[42] (**Supplementary Table 7**). We observe that both CpG islands and CpG methylated sites co-localize strongly with high CpG and GC-content in the human reference (**Supplementary Figure 3**). We do not observe any difference in the distribution of CpG sites hyper or hypo-methylated in human and Great Ape species (data not shown). This shows that these different correlations of mutation density with GC-content are not caused by neither changes in the GC-content of the sequence in Great Apes nor changes in the distribution of hypermutable methylated CpG sites. One strong caveat of this analysis is that methylation was measured from blood samples in all species. It has been shown that germline progenitor cells, specially testis, show increased mutation at methylated CpG sites[43].

### Analysis of non-CpG sites

We divided the dataset and compared independently mutations at non-CpG sites, and CpG>T transitions at CpG sites. We corrected the mutation density at non-CpG sites by the non-CpG fraction of the window passing all our filters. Mutation density at non-CpG sites shows between-datasets correlations similar to the correlations obtained when using mutations in all sites, with slightly stronger correlations between human and NHGAs, and between tumor and all datasets, still maintaining their differences (**Figure 3**). The comparison of mutation density with the CpG content of the window shows a similar distribution as when using all sites, although both human and NHGAs have (as expected), lower CpG content in windows with higher mutation density. This puts NHGAs at a similar level with tumors, with humans being slightly higher, both when using the ranked (**Figure 3**) or unranked (**Supplementary Figure 3**) mutation density. When comparing human, chimpanzee and tumor, we obtain a similar distribution as when using all sites. Using mutations at non-CpG sites, though, mostly removes the diagonal split pattern seen before and instead shows a more horizontal divide (**Supplementary Figure 3**) like in the case of comparing high and low-diversity NHGAs (**Supplementary Figure 2**). Using non-CpG sites we observe still some structure in the GC-content, unlike in the case of NHGAs subspecies, suggesting that non-CpG, GC-rich sites have increased mutation density in humans. Mutations at non-CpG sites in low-diversity NHGA subspecies show weaker correlations with all datasets (as humans do) than high-diversity subspecies

### Analysis of CpG>T transitions

Transitions at CpG sites (xCG>T) when corrected by the size of the window passing all filters is similar to an account of the number of CpG>T mutations in the window. This measure shows extremely high correlations between all the germline datasets (**Figure 3**) but no correlation with tumor. The relationship between mutation density and CpG in the window (using both ranked and unranked data) is identical in the germline datasets (with more mutated windows having more CpG content), while very different in tumor (windows with more CpG>T mutations tend to have low CpG content, and high-CpG content is present in windows with low mutation density). This difference between germline and somatic datasets can be explained by an increased mutation rate of methylated CpG sites in germline cells[43]. The combination of this distribution of uncorrected CpG>T transitions and mutations at non-CpG sites reconstruct the distribution observed when using all mutations together: human and NHGAs have similar distributions in CpG>T transitions, but different in non-CpG mutations, where NHGAs resemble tumor. The sum of both mutation types gives as a result the distribution seen pooling all samples together (**Figure 3 & Supplementary Figure 3**). Humans have very strong correlations of uncorrected CpG>T transitions with all datasets. However, low-diversity NHGA subspecies have much weaker correlations than both humans and high-diversity NHGA subspecies (**Supplementary Table 3**). This suggests an effect on CpG>T sites of human-specific events.

When correcting CpG>T transitions by the number of CpG sites in the window reference, the pattern inverts, and now all datasets, including tumor show extremely high correlations (**Supplementary Figure 3**). All datasets show a very similar distribution between mutation density and CpG-content, with CpG-rich windows having lower mutation density than CpG-poor windows. The differences in the distribution of CpG>T when correcting or not by the number of CpG in the sequence can be explained by CpG-rich windows having more CpG-islands which tend to be unmethylated, and thus, do not spontaneously deaminate. CpG-rich windows have more CpG>T transitions in total number, as there are more CpG bases to mutate, but they have less CpG>T mutation rate than windows with less CpG sites[44]. When comparing the relative mutation rates of CpG>T and non-CpG sites (**Supplementary Figure 3**), we observe that windows in human have a poor correlation between the mutation rate of CpG>T and non-CpG (1kGP R=0.39; 1kGP_69: 0.54; sgdp_50: 0.56), while the correlation is much stronger in NHGAs (chimpanzee R=0.67, gorilla R=0.84), and extremely high in tumors (R=0.94). This suggests that the mutation rates of CpG sites and non-CpG sites are decoupled in the human germline. One possible explanation for this effect is a recent population expansion in human, which caused a higher accumulation of the more clock-like CpG>T transitions. Another possible explanation would be a saturation of CpG>T transitions in Great Ape and tumor genomes, as the number of xCG sites is smaller than any other trinucleotide in the genome and a high number of CpG>T mutations probabilistically lead to repeated mutations in different samples.

### Correlation of CpG>T / non-CpG mutations and genomic features

In a previous analysis, we have described how NHGAs had correlations with certain genomic features more similar to tumor than to the human germline ones. Several of these features, like the histone marks H3K36me1 or H3R2me1 are highly associated with CpG sites[45,46]. As we have seen, the mutation densities at CpG and non-CpG sites vary between datasets.

We analyzed the correlation of the different genomic features with the different types of mutations. Uncorrected CpG>T mutations (which have an extremely high correlation between germline datasets, and no correlation with tumor), show high correlations with all high-CpG-content and open chromatin-associated genomic features in germline datasets (similar in all), and moderate to strong negative correlations in tumor. Using the sequence-corrected CpG>T mutation density (highly correlated in all germline and tumor datasets, and inversely correlated with CpG-content), shows identical strong negative correlations with CpG-associated features. However, when using mutations at non-CpG sites (similar in NHGAs and tumor, and different in human), we observe similar strong correlations of NHGAs and tumor with low-CpG, closed chromatin-associated features, while humans show much weaker correlations in the same direction, albeit stronger than when using all sites (**Supplementary Figure 2**).

Similar as the case of the global mutation density being affected differently by CpG>T and non-CpG mutations, the correlations with genomic features are also affected by both types of mutations. While non-CpG mutations in all datasets correlate with closed chromatin features (strongly in tumor and NHGA, moderately or weakly in humans), CpG>T correlates strongly and positively with open chromatin features in germline datasets. The combination of both effects creates the gradient seen using all variants

(**Figure 2a**) where tumors have a strong correlation with closed-chromatin features, NHGAs have moderate correlations, and humans have very weak, non-existent, or are even correlated with open-chromatin features.

### Recurrent mutations at CpG sites

It has been previously reported that hypermutable CpG sites are prone to mutations appearing independently in different samples[47,48]. Taking into account that the NHGAs mutation rate per year is 1.5x higher than in humans[49], we hypothesized that the mutation density calculation in NHGAs might be biased by recurrent CpG mutations. We analyzed the number of doubleton variants at CpG sites that are shared between populations or subspecies in all germline datasets (**Supplementary Table 8**). When dividing the datasets into a similar number of groups (otherwise, having more groups would lead to more between-group matches), we observe a similar proportion of SNV doubletons at CpG sites that are shared between distant populations. This suggests that the effect of recurrent mutations at CpG sites is similar in all datasets. Thus, the effect of CpG>T saturation must be similar in all germline datasets.

### Human *de novo* mutations

*De novo* mutations offer an unbiased way to assess recurrent CpG>T mutations. We used a dataset of 77k *de novo* mutations in human trios[50] (after filtering). We used the liftOver tool to map the original dataset to the hg19 human reference.

Using all mutations pooled together, human *de novo* mutations present a much strong correlation with population-based human germline datasets, weak with NHGAs germlines, and almost no correlation with tumor datasets (**Supplementary Table 9**). Due to the much smaller number of mutations in the *de novo* dataset, we repeated the correlation analysis using Poison Regression and Negative Binomial Regression, methods better suited to deal with the stochasticity of data based in counts. Both methods showed similar correlations between datasets as the Pearson's correlation.

We divided the human *de novo* mutations into non-CpG and CpG>T mutations. Non-CpG mutations showed similar correlations as all mutations pooled together (moderate with human, weaker with NHGAs, even weaker with tumors). CpG>T human *de* novo mutations showed strong correlations with all the germline datasets, and no correlation with tumor (**Supplementary Table 9**). This suggests that, although we detect differences between human *de novo* mutations and population-based mutations in both humans and NHGAs (and more so in tumor), these differences are not driven by CpG>T mutations, which show similar correlations in the germline of all studied Great Apes. The observed differences in non-CpG sites may still be caused by the small number of mutations of the *de novo* dataset. Further analysis would be needed to assess this effect.

### Analysis of Trinucleotides

Previous studies have show that the mutation rate of a nucleotide is affected by the adjacent sequence context[51]. Trinucleotides are the minimum unit used for sequence context analysis, composed by the mutated nucleotide and the adjacent sites at both the 5' and 3' ends. Combined with 3 possible mutations, this gives a total of 192 possible sequences that, once folded by the complementary sequence, leave 96 possible combinations of sequence and mutation.

One major caveat when comparing mutations between close species is the definition of the ancestral and derived alleles. This is extremely important when analyzing trinucleotides, as misidentifying the ancestral allele will classify a mutation as a completely different trinucleotide, biasing our analysis. We filtered the SNV data in all datasets in order to use only a set of high confidence variants using the following criteria:

- The trinucleotide sequence can be mapped from the human hg19 reference to the chimpanzee reference panTro5 and the gorilla reference gorGor5 using the liftOver tool.
- The trinucleotide sequence must be identical for all 3 alleles in all 3 reference genomes (hg19, panTro5, and gorGor5), accounting by strand changes.
- The trinucleotide sequence cannot overlap the borders of our previously defined window filters (all 3 basepairs must be within the part of the window passing our filters).
- The trinucleotide sequence cannot overlap a multi-nucleotide variant (two or more adjacent SNV variants called in the same sample).
- The trinucleotide cannot overlap fixed SNV in the adjacent +1/-1bp in another species.

- The trinucleotide cannot overlap an InDel in the analyzed species.
- The trinucleotide cannot overlap an InDel with frequency ≥50% in the other two species (for example, in the case of analyzing chimpanzee, remove InDels with AF≥50% in human or gorilla).

The resulting subset of high-confidence variants leaves 47,872,324 SNV in the human 1kGP dataset (81.78% of the original dataset), 10,651,708 SNV (73.38%) in the human sgdp_50 dataset, 26,871,923 SNV (77.61%) in the chimpanzee dataset, 9,813,306 SNV (63.46%) in the gorilla dataset, and 30,756,260 SNV (90.24%) in the tumor dataset. This filtering affects mostly the NHGA datasets, as they are most distant of the human reference used. In addition, the NHGA reference genomes are built using an individual of a given subspecies, causing misidentifications of the ancestral and derived alleles when compared with the human reference. For example, if an SNV is present only in the central chimpanzee subspecies, there is a chance that this SNV was set as the reference allele in the chimpanzee reference genome, when the rest of chimpanzee subspecies carry the same ancestral allele as the human reference.

This 3-sequence filtering affects more CpG sites (85% passing) than non-CpG sites (95% passing) in humans, caused by the hypermutability of CpG sites, where many of them are mutated and annotated as TpG or CpA sites in other references. This sequence bias is found also in NHGAs, although the difference between CpG and non-CpG sites is smaller (**Figure 4, Supplementary Table 10**).

### Biased assessment of CpG sites

Our assessment of mutations at CpG sites is, by definition, biased. CpG>T transitions represent 10.6% of all trinucleotide variants in the 1kGP dataset (chimpanzee: 9.9%; gorilla: 10.3%). Once we apply our 3-sequence filtering, they become 9.5% of all passing sites in the 1kGP dataset (chimpanzee: 9.3%; gorilla: 9.8%). Previous *de novo* mutations in human give CpG>T estimates between ~13%[52], and 16~17%[50,53,54]. This difference could be expected just by the nature of the data used. *De novo* mutations account for new mutations in each individual, while our analysis of segregating sites in population data masks independent mutation events in the same site as a single mutation event. This has been previously shown by the 7~8% of doubleton CpG>T shared between distant populations in the same species (**Supplementary Table 8**). Previous studies have suggested that the mutation rate of CpG>T transitions is identical in the Great Ape lineage [55], and a recent study of de novo mutations in NHGA trios have shown CpG>T represent 17~32% of all mutations in chimpanzee, 14% in gorilla, and 19% in orangutan. This suggests that there are no significant biases in CpG>T mutation rates between human and NHGAs, while there are important differences in the estimates of individual studies[56] or even families[53], while still being similar in all species[55]. In addition to this variation, and the inability of our method to detect recurrent mutations, our choice of analyzing species-exclusive SNV combined with the choice of a human dataset with big sample size (we select species-exclusive SNV in NHGA species comparing against the human dataset 1kGP, with 50x larger sample size than sgdp 50) causes additional biases to all estimates of enrichment in CpG>T transitions. Because all of this, we will ignore all the enrichment results obtained in xCG>T trinucleotides, as we will deem them not conclusive.

### Correlation with mutation signatures

We downloaded the catalog of COSMIC[57] mutation signatures (v3 May 2019) from https://cancer.sanger.ac.uk/cosmic/signatures/SBS/. We correlated all datasets with Signatures SBS1 and SBS5. These signatures have been described by previous studies to reconstruct the mutation pattern of de novo mutations in human[53,58]. We observe very high correlations of all datasets with signature SBS5, with much lower correlations with SBS1 as expected (SBS1 has few important components) (**Supplementary Table 10**). We combined SBS1 and SBS5 in a single signature, applying a correcting factor to each signature equivalent to the relative weight of CpG mutations (SBS1) or non-CpG mutations (SBS5). This resulted in even stronger correlations with the combined signature. It is worth noting that previous studies[53] used a similar approach but used much higher weights for the CpG mutations (25% in their study, 10% in ours). This difference is probably caused by the recurrent nature of CpG>T transitions. These mutations can be seen as recurrent mutations when using population data, while studies of *de novo* mutation can observe them in each individual sample.

### Species-exclusive variants

We analyzed all trinucleotides in the human, chimpanzee, and gorilla datasets, and determined if each mutation is species-exclusive or it is shared between two or more datasets. The majority of shared variants are CpG>T transitions, as well as a smaller number of T>C transitions. An important caveat of this analysis is that the choice of human dataset will have a very big impact. The 1kGP human dataset has 2,504 samples. It has 1,516,059 ACG>T mutations of which 1,125,648 are species-exclusive, and the rest are shared with chimpanzee, gorilla, or both. The human dataset SGDP_50 has 50 samples, 362,142 ACG>T mutations of which 264,804 are species-exclusive. The chimpanzee dataset, when compared with the gorilla and 1kGP human dataset, has 559,482 species-exclusive ACG>T variants, but, when compared with the SGDP_50 human dataset, it has 771,341 species-exclusive ACG>T variants. Gorilla has 184,092 ACG>T when compared with 1kGP, and 254,224 ACG>T when compared with SGDP_50.

On one hand, this shows that the choice of human dataset will bias the results of the analysis of CpG>T transitions in other datasets. But, on the other hand, this suggests that these shared CpG>T variants are repeated variants and not shared variants. These variants are at low frequency (<1%) in the human population, as they are not captured by the SGDP_50 dataset. Only CpG>T trinucleotides are strongly affected by the choice of dataset.

### Conservation of the mutation spectra in primates

We obtained population data from 3 species: the Vervet monkey[59] , Mouse[60], and Pig[61]. We calculated the mutation spectra of each species' population mapped to their own species' reference genome. The mutation spectrum of Vervet monkeys is highly correlated with both signatures SBS1, SBS5, their combination, and with human, chimpanzee and gorilla germlines. Mouse and pig show poorer correlations (**Supplementary Table 10**). This suggests that the mutation spectra are conserved in the primate lineage, ad much more so within the Great Ape lineage.

### Whole-Genome enrichment of trinucleotides

We calculated if there is a significant enrichment of certain trinucleotides in human when compared with other NHGA species. To do this, we used the modified chi-square method described in Harris, eLife 2017[62] and set a p-value threshold of 1e-5 to determine its significance. We calculated also the fold-enrichment of each trinucleotide.

The choice of the human dataset, as previously described, has an effect in CpG>T transitions. When comparing species-exclusive trinucleotides in human and chimpanzee, two CpG>T transitions (ACG>T, and TCG>T) show non-coherent fold enrichments when using either the 1kGP or the SGDP_50 human datasets (more enriched in human when using 1kGP, more enriched in chimpanzee when using SGDP_50). These were the only trinucleotide types affected by the dataset choice. We thus reported as significant only those trinucleotides that showed the same direction of enrichment in both the 1kGP and SGDP_50 datasets, reporting the 1kGP fold-enrichment value (**Figure 4**). Pooling species-exclusive and shared variants, or using only non-singleton variants showed no change in the general enrichment distribution (data not shown).

The top 10% most differently enriched trinucleotides in each of the three comparisons (human – chimpanzee, human – gorilla, chimpanzee – gorilla) were compared against the list of mutation signatures where that trinucleotide appeared at ≥ 5% frequency. Only ATT>A and TTA>A appeared associated with a signature at high frequency (SBS34, 31.6% and 27.2% respectively), although this signature had no associated aetiology yet. The rest of the trinucleotides were associated with signatures in which the trinucleotide has very low frequency or the signature is potentially an artifact (**Supplementary Table 11**).

Our previous analyses have suggested that certain non-CpG mutations in GC-rich windows might have higher mutation density in human than in NHGAs (**Figure 3, Supplementary Figure 3**). We hypothesized that this might be detected at trinucleotide level. Thus, we built a linear regression model comparing the log10 fold-enrichment value as a function of the number of C and G in the trinucleotide's 3bp sequence (triplet). This linear model shows that the number of C/G bases in the sequence has a significant effect in the trinucleotide fold enrichment in the human – chimpanzee comparison (p-value=6.88e-5) (**Supplementary Figure 4**). This significant association holds when including or removing xCG trinucleotides (p-value=3.97e-9) or even when removing TTA>A, the most enriched trinucleotide in chimpanzee (data not shown). This effect is non-significant in human – gorilla comparisons, but significant in chimpanzee – gorilla comparisons. This suggests that GC-rich trinucleotides are more enriched in human than in chimpanzee, but this effect is negligible when comparing human and gorilla. AT-rich trinucleotides are more enriched in chimpanzee than in gorilla.

### Distribution of trinucleotides across the genome (trinucleotide-difference test)

We analyzed the distribution across the genome of trinucleotide variants. This allow us to detect if the human-NHGA-tumor diagonal pattern observed using all variants could be observed when using only a given trinucleotide. To do this, we selected 4,920 1Mbp windows where at least 50% of the window passed both the mappabiliy+cnv+callability filter and the reference genome of human, chimpanzee, and gorilla had the same sequence. We counted the number of mutations of each of the 96 trinucleotides in each window, and divided it by the fraction of the window sequence that has that trinucleotide's corresponding 3-bp reference sequence. This gives a measure of mutations / 1Mbp for each trinucleotide mutation and reference, regardless of sequence content. We calculated the standardized mutation distribution in each of the human, chimpanzee, gorilla, and tumor datasets, and also the ranked distribution.

For each window, we created two distributions: in the first one, we subtracted the ranking in the human and tumor datasets. In the second one, we subtracted the ranking in the NHGA (chimpanzee and gorilla separately) and tumor datasets. These human-tumor and chimpanzee-tumor distributions resemble a normal distribution with mean 0. The standard deviation of these distributions is a magnitude of how similar or different are the tumor and human/NHGA datasets(data not shown). We calculated the Kolmogorov-Smirnov test (R function ks.test) of both human-tumor and chimpanzee-tumor:

$$KS\ test\ (\ (Rank\ Human - Rank\ Tumor)\ , (Rank\ NHGA - Rank\ Tumor)\ )$$

If human and chimpanzee have a similar relationship with tumor (being either both human and NHGAs similar to tumor, or equally different), both the human-tumor and the chimpanzee-tumor distributions would be similar according to the Kolmogorov-Smirnov test. Otherwise, The KS test will mark both distributions as significantly different. The difference between the standard deviations in both distributions can be used as a magnitude of the effect of how different both tumor-subtracted distributions are (**Figure 4**).

In addition to the KS-test p-value, we calculated the partial correlation between human – tumor, and chimpanzee – tumor using the R package ppcor. The partial correlation reports the correlation of a given trinucleotide between human and tumor, once the correlation between human and chimpanzee, and chimpanzee and tumor has been factored out. The combination of the KS-test p-value and the partial correlations show good discriminatory power (**Supplementary Figure 4**). All trinucleotides deemed to be significantly different show a strong clustering, with much stronger partial correlations between NHGAs and tumor than between human and tumor.

We did not take into account the SGDP_50 human dataset for this trinucleotide-difference analysis because it had a smaller number of SNV than 1kGP. This smaller number of SNV, once divided into 4,920 windows, caused a lot of noise and spurious correlations. Nonetheless, the significant results of ACG>T and TCG>T in **Supplementary Figure 4**, again, are probably caused by the effect of different sample size in species-exclusive SNV.

We also modeled the effect that the number of C/G basepairs of the trinucleotide has in the difference of standard deviations results of the trinucleotide-difference test (**Supplementary Figure 4**). We observe no significant association in human-chimpanzee comparisons (**Supplementary Figure 4**), and extremely weak correlations in human-gorilla (p-value=0.036) and chimpanzee-gorilla (p-value=0.008) comparisons (data not shown).

### Effect of cancer mutational signatures in individual tumor types (signature-difference test)

We ran the trinucleotide-difference test using each individual tumor type instead of pooling all tumor types together. This shows almost none new significant association between individual tumor types and NHGAs that were not detected using all tumors combined. However, some individual tumor types tended to recapitulate the results of using all tumors (**Supplementary Table 12**).

Each individual tumor type is affected in different proportions by the different cancer mutation signatures. We tested if the results of the trinucleotide-difference test in individual tumor types were associated with specific mutation signatures. We downloaded mutation load data for each signature and each tumor type from the PCAWG project[63], and calculated the number of mutations/Mbp per sample associated to each mutation signature in each individual tumor type. For each cancer signature (65 signatures) and trinucleotide (96 trinucleotides) combination, we built a linear regression where the trinucleotide-difference test effect size (the difference between the distributions' standard deviations) of the trinucleotide in each tumor type was dependent on the average mutation load of the signature in that tumor type. We obtained the p-value associated with the mutation load in this linear model, and run a chi-squared test counting the number of trinucleotides significant (p-value <0.05) in both the trinucleotide-difference test and the linear model. Only two signatures had significant p-value after multiple correction (p-value <10e-4): Signatures SBS5 and SBS40 (**Figure 4**). Both are “flat” signatures affecting multiple mutation types. They affect a high number of samples in most if not all tumor types, and have been associated with the age of the patient sample[57,63]. Signature SBS5 has been associated previously with healthy somatic cell mutation accumulation and *de novo* germline mutations[53,64,65]. Notoriously, signature SBS1 is also associated with the sample’s age, but it is not found significant, mostly because signature SBS1 is composed mainly of CpG>T transitions and we detect a similar distribution of those in humans and NHGA in the trinucleotide-difference test.

We repeated this signature-difference test using more complex linear regression models using additional variables: the total number of mutations associated with a signature in all samples in the tumor type, the total number of mutations in the tumor type, and the number of samples in the tumor type. Combinations of these variables with the mutation load of a signature were not informative nor significant in the linear regressions, as these variables are not independent (the total number of signature mutations in a tumor type is dependent on the signature’s mutation load and the number of samples; the total number of mutations is dependent on the number of samples and constant across signatures, etc.). Using only the signature’s mutation load was the best predictor of the result of the trinucleotide-difference test across tumor types.

### Supplementary Tables legends

**Supplementary Table 1:**

Partial correlations. **a)** Partial Pearson's correlations comparing 1kGP-chimpanzee-tumor; **b)** comparing 1kGP-gorilla-tumor.

**Supplementary Table 2:**

Correlations using different filters. **a)** Using different thresholds for the minimum fraction of the 1Mbp windows passing our filtering; **b)** using different window sizes with a minimum window fraction of 50%; **c)** mapping data to the chimpanzee panTro5 reference; **d)** using archaic hominids.

**Supplementary Table 3:**

Correlations of subspecies. **a)** Pearson's correlation R of all pairwise correlations between diverse datasets using all sites. NS: non-significant: p-value > 0.05. 1kGP: 1000 Genomes Project 2,504 samples; 1kGP 69: 1000 Genomes Project 69 samples; 1kGP 43: 1000 Genomes Project 43 samples; ICGC: release of 1,823 germline samples from the PanCancer Analysis of Whole Genomes; SGDP 279: release of 279 samples from Simons Genome Diversity Project; SGDP 50: our calling of 50 human samples from SGDP; Africa 26: Subset of SGDP 50 with 26 samples of African origin; Not Africa 24: Subset of SGDP 50 with 24 samples of non-African origin. Chimpanzee: our calling of 69 chimpanzee; pp: *Pan paniscus* (bonobo) 10 samples; ptv: *Pan troglodytes verus* 12 samples; pte: *Pan troglodytes ellioti* 10 samples; pts: *Pan troglodytes schweinfurthii* 19 samples; ptt: *Pan troglodytes troglodytes* 18 samples; ptt 5: subset of 5 samples of ptt. Gorilla: our calling of 43 gorilla samples; gbb: *Gorilla* *beringei beringei* 7 samples; gbg: *Gorilla beringei graueri* 8 samples; ggg: *Gorilla gorilla gorilla* 27 samples; ggg 5: subset of 5 samples of ggg. Tumor: release of 2,583 human tumor samples from PCAWG. Window ratio: fraction of the window passing filters; GC-content of the window; CpG dinucleotide content of the window. **b)** Using mutations at non-CpG sites; **c)** using CpG>T transitions

**Supplementary Table 4:**

Correlations of tumor types. Pearson's correlation R between human, chimpanzee, or gorilla standardized mutation density in 1Mbp windows, and the mutation density of tumors of different histologies.

**Supplementary Table 5:**

Allele frequency correlations. **a)** Pearson's correlation R between different species using only variants of a certain frequency; **b)** correlation between variants at different frequencies within each dataset. **c)** correlation between species when using only variants exclusive from a species; **d)** correlation between variants exclusive in one species and variants shared between pairs of human-Great Ape species; **e)** correlation between human-chimpanzee divergence or human-gorilla divergence and all segregating sites in each dataset.

**Supplementary Table 6:**

InDel correlations **a)** Pearson's correlation R between SNV mutation density and InDel mutation density in the different datasets; **b)** InDel mutation density between datasets.

**Supplementary Table 7:**

Source of the different genomic features used.

**Supplementary Table 8:**

Non-CpG / CpG>T fraction. **a)** Fraction of CpG and non-CpG sites with respect of the total variants, the non-singleton variants, and the average number of heterozygous sites per sample. **b)** Fraction of doubletons at CpG sites shared between samples of different subspecies/populations using all mutations at CpG or exclusively CpG>T transitions.

**Supplementary Table 9:**

Correlations with *de novo* mutations. **a)** correlation of the standardized mutation distribution between 77,000 human *de novo* mutations from Carlson 2018 compared with other datasets, using all variants; **b)** using only SNV at non-CpG sites; **c)** using only CpG>T transitions.

**Supplementary Table 10:**

Mutation spectra correlations. **a)** Fraction of trinucleotides at non-CpG and CpG sites passing the 3-references filter; **b)** correlation of the species' mutation spectrum with mutation signatures; **c)** pairwise correlations of species' mutation spectra.

**Supplementary Table 11:**

Enriched trinucleotides presence in signatures. Frequency at which appear in different tumor mutation signatures of the COSMIC catalog, of trinucleotides that are among the top 10% most enriched in human-chimpanzee, human-gorilla, and chimpanzee-gorilla comparisons.

**Supplementary Table 12:**

Trinucleotide-difference test in tumor types. Difference between the standard deviations of the distributions in the trinucleotide-difference test using individual tumor types comparing **a)** 1kGP-chimpanzee, and **b)** 1kGP-gorilla. Only significant (p-value <10e-5) results are shown.

### Supplementary Figures

**Supplementary Figure 1:**

Robustness of the results. **a)** distribution and 2D-density of the unranked mutation density in pairwise comparisons between datasets; **b)** density and distribution of the fraction of the window passing our filters, setting thresholds at 50%, 60%, 70%, and 80%; **c)** distribution of the ranked mutation density using 10Mbp, 1Mbp, 100kbp, and 10kbp window sizes; **d)** distribution of the unranked mutation density when using only windows with ≥25% difference in ranking in both datasets; **e)** comparison of the ranked mutation density between chimpanzee and gorilla; **f)** between NHGA and different human datasets; **g)** between human and chimpanzee mapped to the chimpanzee reference genome, showing tumor and GC-content; **h)** comparison of 3 archaic samples with modern humans and NHGA. The numbers above the figure denote the Mann-Whitney U-test p-value dividing the data by the diagonal.

Suppl. Figure 1 Robustness of the results

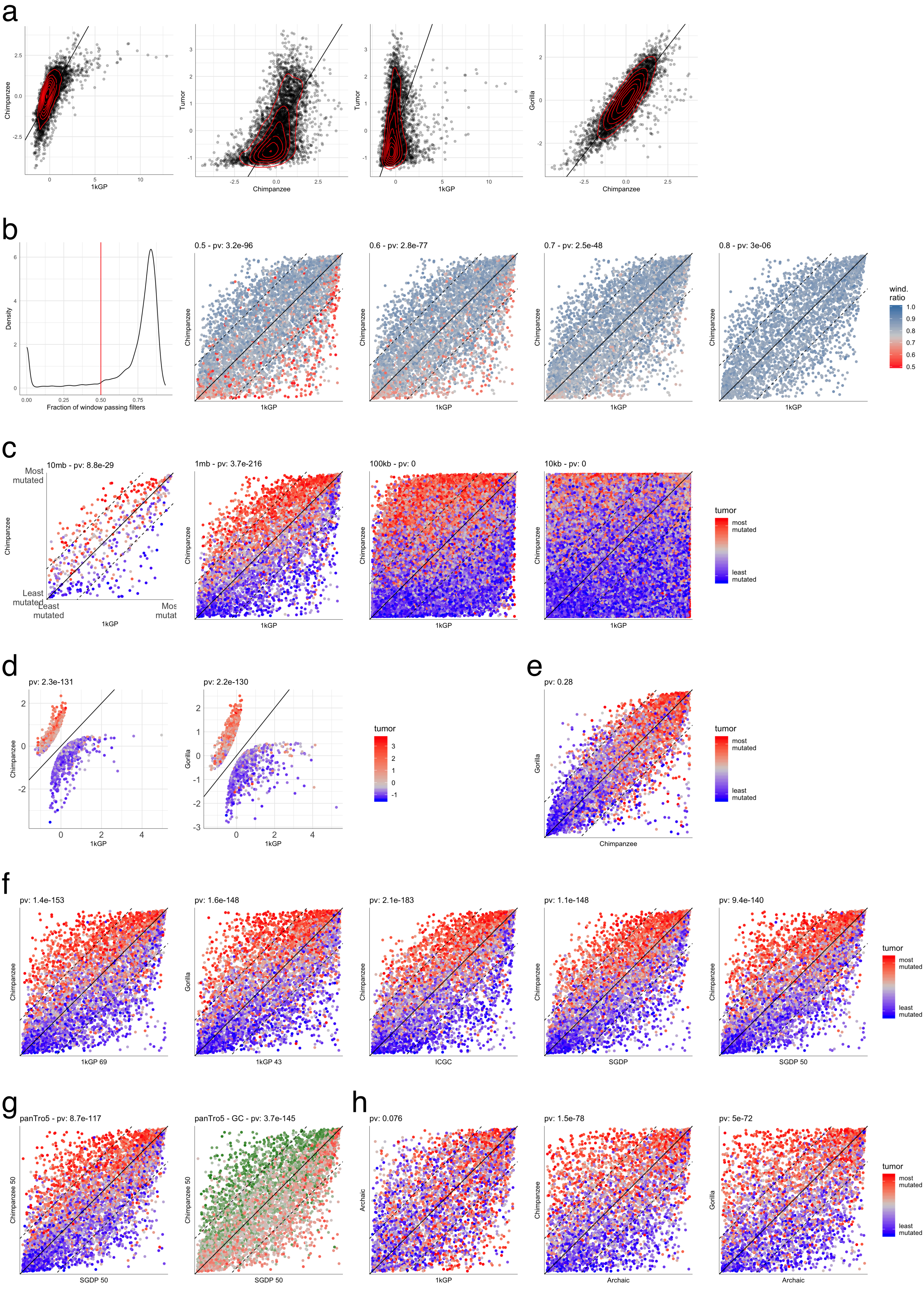

**Supplementary Figure 2:**

Subspecies. Comparisons of the ranked mutation density between human and NHGA subspecies. For a list of subspecies names, check Supplementary Table 3. Human XX denotes a subset of 1kGP samples of African origin (to maximize human diversity) of the same XX sample size as the NHGA subspecies. **a)** chimpanzee subspecies; **b)** gorilla subspecies; **c)** low-diversity (x-axis) vs high-diversity (y-axis) subspecies of the same species; **d)** low-diversity (x-axis) vs high-diversity (y-axis) subspecies of different Great Ape species; **e)** low-diversity chimpanzee and gorillas; **f)** high-diversity chimpanzee and gorillas. **g)** Correlation between genomic and epigenomic features and mutations at non-CpG sites; **h)** CpG>T transitions (not correcting by the sequence context).

Suppl. Figure 2 NHGA subspecies and genomic features

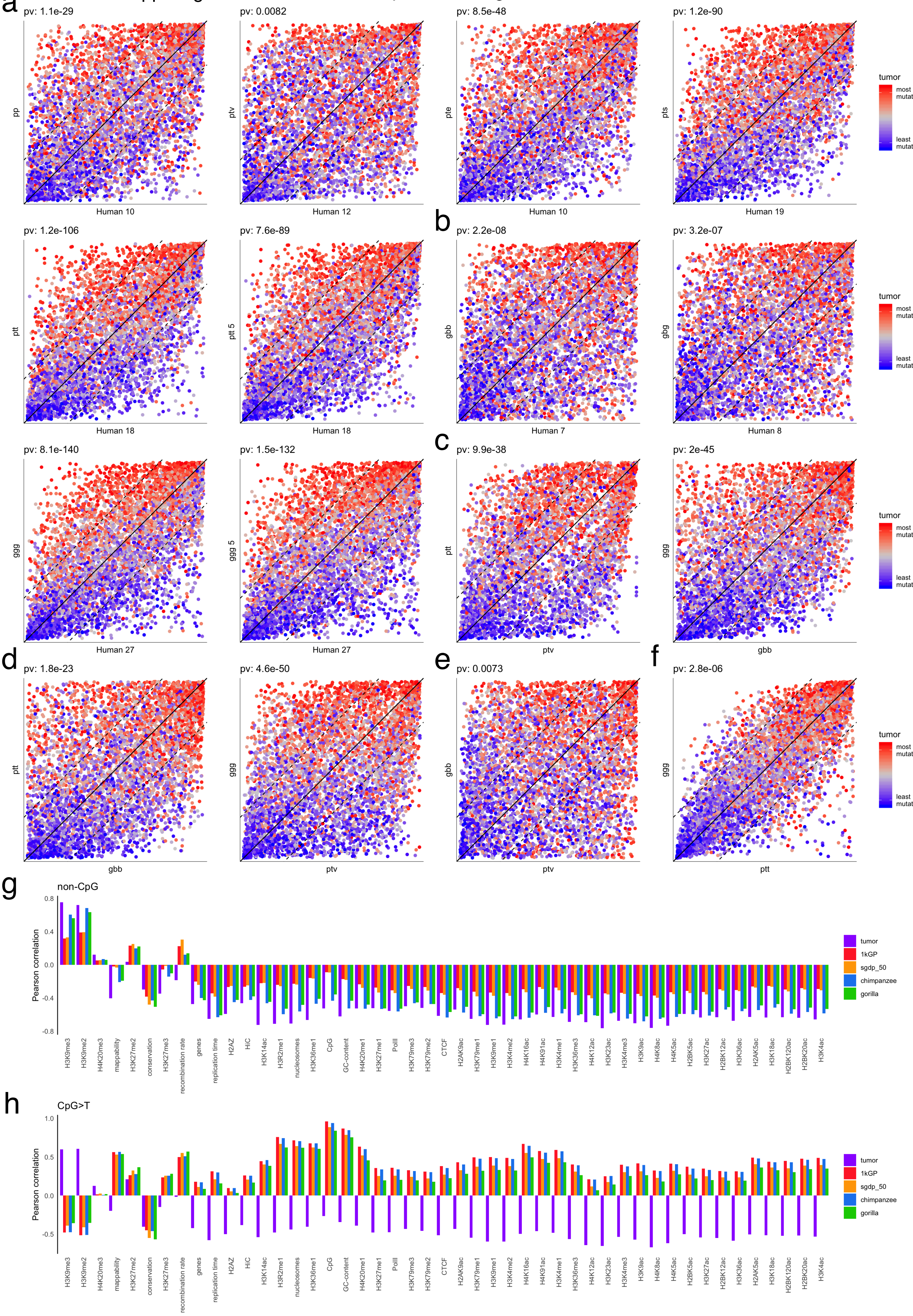

**Supplementary Figure 3:**

GC-content. **a)** distribution of exons, GC-content, CpG-content, CpG-islands, and methylated CpG sites; **b)** distribution of CpG-content comparing a low-diversity chimpanzee (x-axis) and a high-diversity gorilla (y-axis); **c)** loess smooth curves showing the association between ranked mutation density and CpG-content in NHGA subspecies; **d)** association between GC-content in the mappings to the human and chimpanzee reference using all variants; **e)** association between CpG-content and the unranked mutation density (outliers were capped at ±4 ), using all sites, mutations at non-CpG sites, CpG>T transitions, and CpG>T transitions correcting by the number of CpG sites in the sequence; **f)** ranked distribution of non-CpG mutations in human, chimpanzee, tumor, and GC-content; **g)** ranked distribution of CpG>T transitions (correcting by the fraction of CpG sites in the sequence) in human, chimpanzee, tumor, and GC-content; **h)** correlation and distribution between non-CpG and CpG>T (corrected) mutation rate in each window within the same species; **i)** loess smooth curves showing the association between ranked mutation density and recombination rates in different species. The black lines denote the exact subspecies from which the recombination was calculated.

Suppl. Figure 3 CpG–content and recombination

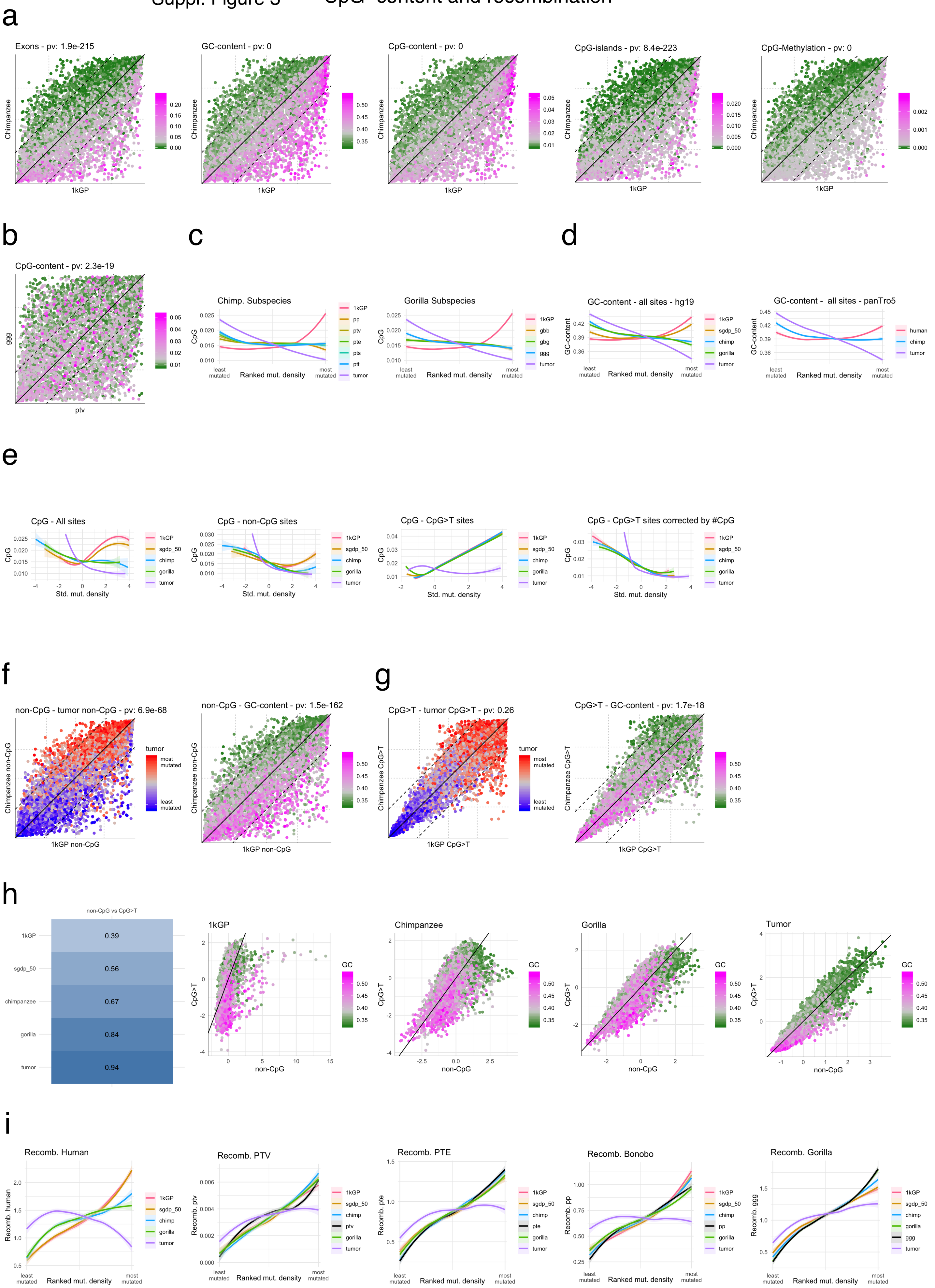

**Supplementary Figure 4:**

Trinucleotides. **a)** mutation spectrum of the 1kGP dataset. The colors denote the trinucleotides failing our 3-reference filter, shared between 2 or more species, or species-exclusive; **b)** in chimpanzee; **c)** in gorilla; **d)** fold-enrichment in 1kGP-chimpanzee vs 1kGP-gorilla (left), and 1kGP-chimpanzee vs chimpanzee-gorilla (right). Labels denote the top 10% significant and most enriched trinucleotides in each axis; **e)** linear regression model test showing a significant association between the number of C and G bases within a trinucleotide and the fold-enrichment in 1kGP-chimpanzee comparisons; **f)** trinucleotides-difference test comparing 1kGP-chimpanzee (left) and 1kGP-gorilla (right). The axis denote the partial correlation 1kGP-tumor (controlling by NHGA) and NHGA-tumor (controlling by 1kGP). Labels and filled circles denote significant trinucleotides (p-value <10e-5); **g)** linear regression model showing no significant association between the trinucleotide-difference test effect size and the number of C and G bases in a trinucleotide, in the 1kGP-chimpanzee comparison.

Suppl. Figure 4 Trinucleotides

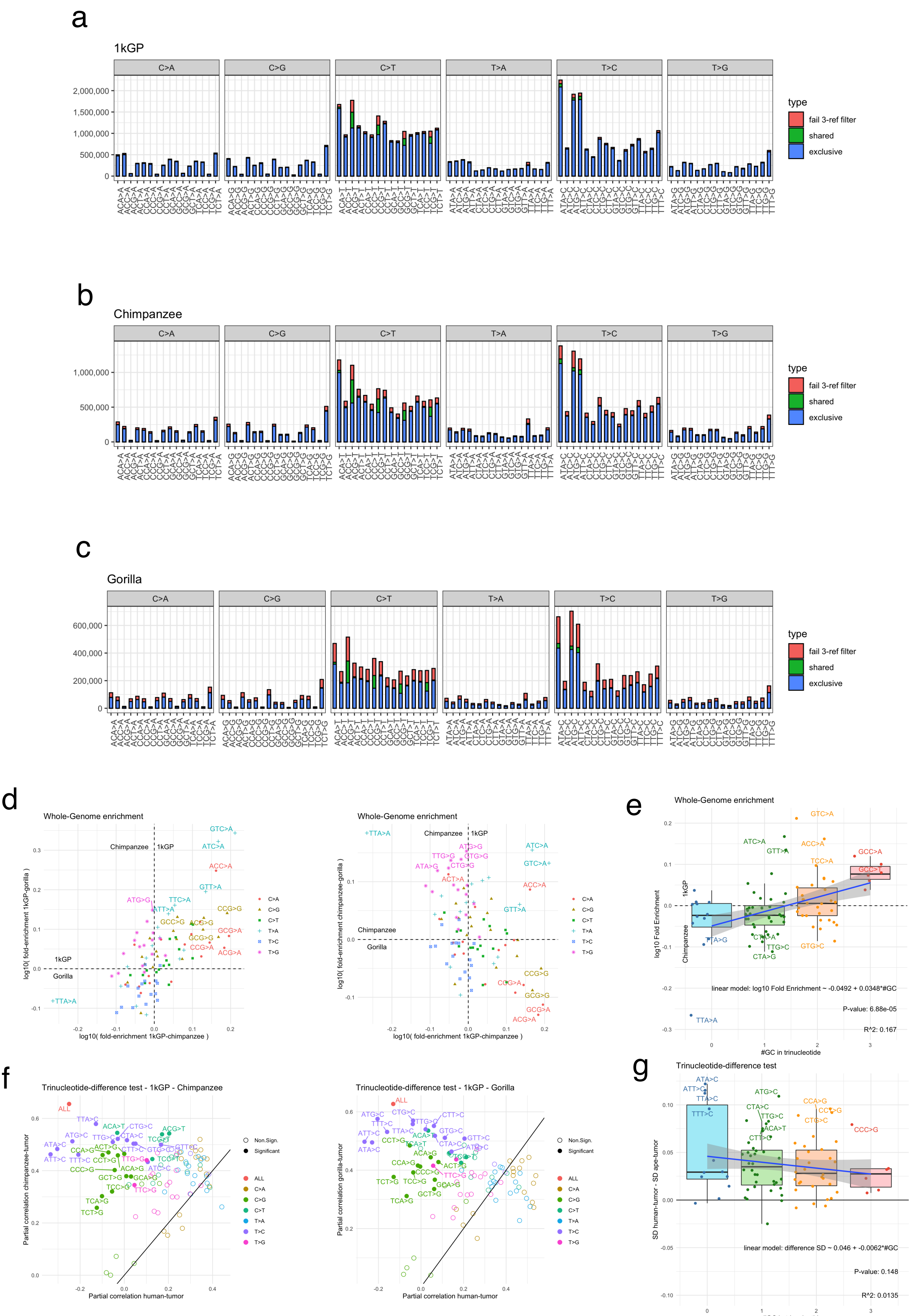
