## Supplementary Tables for "Mutation distribution density in tumors reconstructs human’s lost diversity"

**Supplementary Table 1**

**a)**

Partial Correlations (Pearson's R) - 1kGP-chimpanzee-tumor

| | **1kGP** | **Chimpanzee** | **Tumor** |
|---|---|---|---|
| **1kGP** | - | - | - |
| **Chimpanzee** | 0.67 | - | - |
| **Tumor** | -0.3 | 0.59 | - |

**b)**

Partial Correlations (Pearson's R) - 1kGP-gorilla-tumor

| | **1kGP** | **Gorilla** | **Tumor** |
|---|---|---|---|
| **1kGP** | - | - | - |
| **Gorilla** | 0.54 | - | - |
| **Tumor** | -0.21 | 0.59 | - |

**Supplementary Table 2**

**a)**

Window fraction passing filters

| | 0.5 | 0.6 | 0.7 | 0.8 |
|---|---|---|---|---|
| **Number of windows** | 5040 | 4814 | 4397 | 3037 |
| **human - chimpanzee** | 0.65 | 0.66 | 0.67 | 0.68 |
| **human - gorilla** | 0.53 | 0.54 | 0.55 | 0.54 |
| **chimpanzee - gorilla** | 0.84 | 0.85 | 0.84 | 0.81 |
| **human - tumor** | 0.16 | 0.18 | 0.18 | 0.15 |
| **chimpanzee - tumor** | 0.55 | 0.55 | 0.54 | 0.49 |
| **gorilla - tumor** | 0.58 | 0.58 | 0.57 | 0.52 |

**b)**

Window size

| | 10 Mbp | 1 Mbp | 100 kbp | 10 kbp |
|---|---|---|---|---|
| **human - chimpanzee** | 0.69 | 0.65 | 0.59 | 0.44 |
| **human - gorilla** | 0.54 | 0.53 | 0.49 | 0.36 |
| **chimpanzee - gorilla** | 0.86 | 0.84 | 0.71 | 0.48 |
| **human - tumor** | 0.05 | 0.16 | 0.19 | 0.19 |
| **chimpanzee - tumor** | 0.47 | 0.55 | 0.5 | 0.37 |
| **gorilla - tumor** | 0.53 | 0.58 | 0.5 | 0.37 |

**c)**

PanTro5 mappings

| | sgdp 50 | Chimpanzee 50 | Tumor |
|---|---|---|---|
| **sgdp 50** | - | - | - |
| **Chimpanzee 50** | 0.77 | - | - |
| **Tumor** | 0.25 | 0.52 | - |

**d)**

Archaic hominids

| | 1kGP | archaic | Chimpanzee | Gorilla | Tumor |
|---|---|---|---|---|---|
| **1kGP** | - | - | - | - | - |
| **archaic** | 0.67 | - | - | - | - |
| **Chimpanzee** | 0.66 | 0.58 | - | - | - |
| **Gorilla** | 0.55 | 0.55 | 0.84 | - | - |
| **Tumor** | 0.18 | 0.22 | 0.53 | 0.56 | - |

**Supplementary Table 3**

**a)**
All sites

| | 1kGP | 1kGP 69 | 1kGP 43 | ICGC | SGDP 279 | SGDP 50 | African 26 | Non-African 24 | Chimpanzee | pp | ptv | pte | pts | ptt | ptt 5 | Gorilla | gbb | gbg | ggg | ggg 5 | Tumor | Window ratio | GC-content | CpG |
|---|---|---|---|---|---|---|---|---|---|---|---|---|---|---|---|---|---|---|---|---|---|---|---|---|
| **1kGP** | - | - | - | - | - | - | - | - | - | - | - | - | - | - | - | - | - | - | - | - | - | - | - | - |
| **1kGP 69** | 0.9 | - | - | - | - | - | - | - | - | - | - | - | - | - | - | - | - | - | - | - | - | - | - | - |
| **1kGP 43** | 0.88 | 0.99 | - | - | - | - | - | - | - | - | - | - | - | - | - | - | - | - | - | - | - | - | - | - |
| **ICGC** | 0.99 | 0.91 | 0.89 | - | - | - | - | - | - | - | - | - | - | - | - | - | - | - | - | - | - | - | - | - |
| **SGDP 279** | 0.94 | 0.97 | 0.96 | 0.95 | - | - | - | - | - | - | - | - | - | - | - | - | - | - | - | - | - | - | - | - |
| **SGDP 50** | 0.91 | 0.97 | 0.96 | 0.91 | 0.97 | - | - | - | - | - | - | - | - | - | - | - | - | - | - | - | - | - | - | - |
| **African 26** | 0.89 | 0.96 | 0.96 | 0.89 | 0.95 | 0.99 | - | - | - | - | - | - | - | - | - | - | - | - | - | - | - | - | - | - |
| **Non-African 24** | 0.79 | 0.92 | 0.92 | 0.81 | 0.89 | 0.92 | 0.89 | - | - | - | - | - | - | - | - | - | - | - | - | - | - | - | - | - |
| **Chimpanzee** | 0.65 | 0.69 | 0.69 | 0.64 | 0.7 | 0.74 | 0.74 | 0.65 | - | - | - | - | - | - | - | - | - | - | - | - | - | - | - | - |
| **pp** | 0.48 | 0.57 | 0.57 | 0.49 | 0.56 | 0.59 | 0.59 | 0.55 | 0.76 | - | - | - | - | - | - | - | - | - | - | - | - | - | - | - |
| **ptv** | 0.45 | 0.5 | 0.5 | 0.46 | 0.5 | 0.51 | 0.51 | 0.48 | 0.6 | 0.46 | - | - | - | - | - | - | - | - | - | - | - | - | - | - |
| **pte** | 0.51 | 0.61 | 0.61 | 0.52 | 0.6 | 0.63 | 0.63 | 0.59 | 0.82 | 0.61 | 0.53 | - | - | - | - | - | - | - | - | - | - | - | - | - |
| **pts** | 0.57 | 0.66 | 0.66 | 0.58 | 0.65 | 0.69 | 0.7 | 0.63 | 0.92 | 0.67 | 0.57 | 0.81 | - | - | - | - | - | - | - | - | - | - | - | - |
| **ptt** | 0.6 | 0.66 | 0.66 | 0.6 | 0.66 | 0.71 | 0.71 | 0.63 | 0.96 | 0.69 | 0.58 | 0.82 | 0.92 | - | - | - | - | - | - | - | - | - | - | - |
| **ptt 5** | 0.58 | 0.66 | 0.66 | 0.58 | 0.65 | 0.69 | 0.7 | 0.63 | 0.93 | 0.67 | 0.57 | 0.81 | 0.91 | 0.96 | - | - | - | - | - | - | - | - | - | - |
| **Gorilla** | 0.53 | 0.64 | 0.64 | 0.53 | 0.63 | 0.68 | 0.68 | 0.62 | 0.84 | 0.66 | 0.52 | 0.73 | 0.8 | 0.82 | 0.8 | - | - | - | - | - | - | - | - | - |
| **gbb** | 0.32 | 0.4 | 0.41 | 0.32 | 0.39 | 0.42 | 0.43 | 0.4 | 0.53 | 0.43 | 0.32 | 0.47 | 0.5 | 0.52 | 0.5 | 0.64 | - | - | - | - | - | - | - | - |
| **gbg** | 0.37 | 0.45 | 0.46 | 0.37 | 0.44 | 0.47 | 0.47 | 0.45 | 0.55 | 0.42 | 0.39 | 0.5 | 0.54 | 0.54 | 0.53 | 0.67 | 0.5 | - | - | - | - | - | - | - |
| **ggg** | 0.51 | 0.62 | 0.62 | 0.51 | 0.61 | 0.66 | 0.67 | 0.61 | 0.83 | 0.66 | 0.51 | 0.72 | 0.79 | 0.82 | 0.79 | 0.99 | 0.59 | 0.62 | - | - | - | - | - | - |
| **ggg 5** | 0.46 | 0.58 | 0.58 | 0.46 | 0.56 | 0.62 | 0.63 | 0.57 | 0.81 | 0.64 | 0.49 | 0.7 | 0.77 | 0.79 | 0.77 | 0.96 | 0.58 | 0.6 | 0.97 | - | - | - | - | - |
| **Tumor** | 0.16 | 0.22 | 0.22 | 0.18 | 0.25 | 0.26 | 0.27 | 0.22 | 0.55 | 0.42 | 0.32 | 0.46 | 0.5 | 0.52 | 0.5 | 0.58 | 0.38 | 0.37 | 0.58 | 0.59 | - | - | - | - |
| **Window ratio** | NS | 0.03 | 0.03 | NS | NS | 0.08 | 0.08 | 0.06 | 0.32 | 0.22 | 0.12 | 0.25 | 0.29 | 0.31 | 0.28 | 0.31 | 0.18 | 0.18 | 0.32 | 0.34 | 0.36 | - | - | - |
| **GC-content** | 0.25 | 0.15 | 0.14 | 0.23 | 0.15 | 0.1 | 0.08 | 0.08 | -0.24 | -0.21 | -0.13 | -0.19 | -0.22 | -0.25 | -0.23 | -0.24 | -0.18 | -0.15 | -0.25 | -0.3 | -0.59 | -0.42 | - | - |
| **CpG** | 0.37 | 0.25 | 0.24 | 0.36 | 0.28 | 0.2 | 0.19 | 0.17 | -0.12 | -0.12 | -0.04 | -0.1 | -0.12 | -0.13 | -0.12 | -0.15 | -0.12 | -0.1 | -0.16 | -0.23 | -0.49 | -0.5 | 0.91 | - |

**b)**
non-CpG

| | 1kGP | 1kGP 69 | sgdp 50 | human de novo | Chimpanzee | pp | ptv | pte | pts | ptt | ptt 5 | Gorilla | gbb | gbg | ggg | ggg 5 | Tumor |
|---|---|---|---|---|---|---|---|---|---|---|---|---|---|---|---|---|---|
| **1kGP** | - | - | - | - | - | - | - | - | - | - | - | - | - | - | - | - | - |
| **1kGP 69** | 0.9 | - | - | - | - | - | - | - | - | - | - | - | - | - | - | - | - |
| **sgdp 50** | 0.92 | 0.96 | - | - | - | - | - | - | - | - | - | - | - | - | - | - | - |
| **human de novo** | 0.43 | 0.34 | 0.35 | - | - | - | - | - | - | - | - | - | - | - | - | - | - |
| **Chimpanzee** | 0.7 | 0.68 | 0.74 | 0.26 | - | - | - | - | - | - | - | - | - | - | - | - | - |
| **pp** | 0.56 | 0.59 | 0.62 | 0.21 | 0.8 | - | - | - | - | - | - | - | - | - | - | - | - |
| **ptv** | 0.52 | 0.53 | 0.55 | 0.2 | 0.64 | 0.52 | - | - | - | - | - | - | - | - | - | - | - |
| **pte** | 0.58 | 0.63 | 0.65 | 0.22 | 0.84 | 0.66 | 0.58 | - | - | - | - | - | - | - | - | - | - |
| **pts** | 0.64 | 0.67 | 0.71 | 0.24 | 0.94 | 0.72 | 0.62 | 0.83 | - | - | - | - | - | - | - | - | - |
| **ptt** | 0.66 | 0.67 | 0.72 | 0.24 | 0.97 | 0.74 | 0.62 | 0.84 | 0.94 | - | - | - | - | - | - | - | - |
| **ptt 5** | 0.64 | 0.67 | 0.71 | 0.23 | 0.94 | 0.72 | 0.62 | 0.84 | 0.92 | 0.97 | - | - | - | - | - | - | - |
| **Gorilla** | 0.59 | 0.64 | 0.69 | 0.21 | 0.86 | 0.7 | 0.57 | 0.76 | 0.82 | 0.84 | 0.82 | - | - | - | - | - | - |
| **gbb** | 0.38 | 0.43 | 0.45 | 0.12 | 0.56 | 0.47 | 0.36 | 0.51 | 0.53 | 0.54 | 0.53 | 0.66 | - | - | - | - | - |
| **gbg** | 0.42 | 0.48 | 0.5 | 0.15 | 0.58 | 0.46 | 0.43 | 0.53 | 0.57 | 0.57 | 0.56 | 0.69 | 0.52 | - | - | - | - |
| **ggg** | 0.58 | 0.63 | 0.68 | 0.21 | 0.85 | 0.7 | 0.56 | 0.75 | 0.82 | 0.84 | 0.82 | 0.99 | 0.61 | 0.64 | - | - | - |
| **ggg 5** | 0.56 | 0.61 | 0.66 | 0.2 | 0.84 | 0.69 | 0.54 | 0.74 | 0.8 | 0.82 | 0.8 | 0.97 | 0.6 | 0.63 | 0.98 | - | - |
| **Tumor** | 0.38 | 0.34 | 0.39 | 0.16 | 0.66 | 0.53 | 0.41 | 0.55 | 0.6 | 0.62 | 0.6 | 0.66 | 0.43 | 0.44 | 0.66 | 0.66 | - |

**c)**
CpG>T

| | 1kGP | 1kGP 69 | sgdp 50 | human de novo | Chimpanzee | pp | ptv | pte | pts | ptt | ptt 5 | Gorilla | gbb | gbg | ggg | ggg 5 | Tumor |
|---|---|---|---|---|---|---|---|---|---|---|---|---|---|---|---|---|---|
| **1kGP** | - | - | - | - | - | - | - | - | - | - | - | - | - | - | - | - | - |
| **1kGP 69** | 0.97 | - | - | - | - | - | - | - | - | - | - | - | - | - | - | - | - |
| **sgdp 50** | 0.97 | 0.99 | - | - | - | - | - | - | - | - | - | - | - | - | - | - | - |
| **human de novo** | 0.61 | 0.58 | 0.59 | - | - | - | - | - | - | - | - | - | - | - | - | - | - |
| **Chimpanzee** | 0.98 | 0.95 | 0.96 | 0.6 | - | - | - | - | - | - | - | - | - | - | - | - | - |
| **pp** | 0.91 | 0.91 | 0.91 | 0.54 | 0.94 | - | - | - | - | - | - | - | - | - | - | - | - |
| **ptv** | 0.85 | 0.83 | 0.84 | 0.52 | 0.86 | 0.8 | - | - | - | - | - | - | - | - | - | - | - |
| **pte** | 0.91 | 0.92 | 0.92 | 0.55 | 0.94 | 0.88 | 0.82 | - | - | - | - | - | - | - | - | - | - |
| **pts** | 0.95 | 0.94 | 0.94 | 0.57 | 0.97 | 0.91 | 0.84 | 0.94 | - | - | - | - | - | - | - | - | - |
| **ptt** | 0.96 | 0.95 | 0.95 | 0.58 | 0.99 | 0.92 | 0.85 | 0.95 | 0.98 | - | - | - | - | - | - | - | - |
| **ptt 5** | 0.94 | 0.94 | 0.94 | 0.57 | 0.97 | 0.91 | 0.85 | 0.94 | 0.97 | 0.99 | - | - | - | - | - | - | - |
| **Gorilla** | 0.93 | 0.94 | 0.95 | 0.56 | 0.94 | 0.9 | 0.82 | 0.91 | 0.93 | 0.94 | 0.93 | - | - | - | - | - | - |
| **gbb** | 0.67 | 0.71 | 0.71 | 0.41 | 0.69 | 0.66 | 0.59 | 0.69 | 0.7 | 0.7 | 0.69 | 0.77 | - | - | - | - | - |
| **gbg** | 0.76 | 0.78 | 0.79 | 0.46 | 0.77 | 0.74 | 0.69 | 0.75 | 0.77 | 0.77 | 0.77 | 0.84 | 0.7 | - | - | - | - |
| **ggg** | 0.91 | 0.94 | 0.94 | 0.55 | 0.93 | 0.9 | 0.81 | 0.9 | 0.92 | 0.93 | 0.93 | 1 | 0.75 | 0.81 | - | - | - |
| **ggg 5** | 0.87 | 0.9 | 0.91 | 0.52 | 0.89 | 0.87 | 0.77 | 0.87 | 0.89 | 0.9 | 0.89 | 0.97 | 0.74 | 0.8 | 0.98 | - | - |
| **Tumor** | -0.13 | -0.04 | -0.05 | -0.09 | -0.14 | -0.09 | -0.09 | -0.06 | -0.08 | -0.09 | -0.08 | -0.01 | 0.06 | 0.01 | 0 | 0.04 | - |

**Supplementary Table 4**

| Tumor type | # samples | # SNV | Pearson's R human | Pearson's R chimpanzee | Pearson's R gorilla | Mann-Whitney U test p-value |
|---|---|---|---|---|---|---|
| All | 2,583 | 34,911,808 | 0.16 | 0.55 | 0.58 | 3.7E-216 |
| Skin-Melanoma | 107 | 9,601,997 | 0.16 | 0.46 | 0.49 | 1.8E-118 |
| ColoRect-AdenoCA | 52 | 7,281,777 | 0.08 | 0.5 | 0.52 | 1.5E-294 |
| Liver-HCC | 315 | 3,076,372 | 0.1 | 0.49 | 0.51 | 1.7E-221 |
| Eso-AdenoCa | 97 | 2,136,133 | 0.17 | 0.51 | 0.53 | 2.5E-186 |
| Lung-SCC | 47 | 1,616,150 | 0.09 | 0.46 | 0.5 | 1.1E-200 |
| Stomach-AdenoCA | 68 | 1,263,860 | 0.18 | 0.52 | 0.53 | 1.1E-180 |
| Panc-AdenoCA | 232 | 1,183,031 | 0.21 | 0.49 | 0.52 | 6.5E-123 |
| Breast-AdenoCa | 195 | 1,063,688 | 0.17 | 0.39 | 0.4 | 5E-86 |
| Lung-AdenoCA | 37 | 1,009,893 | 0.18 | 0.52 | 0.56 | 4.3E-156 |
| Lymph-BNHL | 105 | 985,991 | 0.13 | 0.41 | 0.42 | 2.2E-215 |
| Ovary-AdenoCA | 110 | 758,465 | 0.16 | 0.45 | 0.47 | 4.5E-107 |
| Uterus-AdenoCA | 44 | 736,102 | 0.13 | 0.45 | 0.47 | 1.9E-191 |
| Kidney-RCC | 143 | 733,510 | 0.14 | 0.43 | 0.46 | 6.2E-97 |
| Head-SCC | 56 | 681,568 | 0.11 | 0.47 | 0.5 | 1.6E-195 |
| Prost-AdenoCA | 199 | 465,385 | 0.17 | 0.46 | 0.47 | 2.8E-132 |
| Bladder-TCC | 23 | 391,265 | 0.14 | 0.39 | 0.4 | 5.5E-80 |
| CNS-GBM | 39 | 377,586 | 0.2 | 0.38 | 0.42 | 3E-15 |
| Biliary-AdenoCA | 33 | 373,854 | 0.18 | 0.49 | 0.5 | 3.1E-150 |
| Panc-Endocrine | 81 | 192,608 | 0.19 | 0.47 | 0.49 | 7.6E-95 |
| Lymph-CLL | 90 | 171,011 | 0.13 | 0.45 | 0.46 | 1.7E-174 |
| CNS-Medullo | 141 | 150,130 | 0.15 | 0.42 | 0.44 | 1.9E-81 |
| Bone-Leiomyo | 34 | 140,410 | 0.13 | 0.34 | 0.38 | 5E-102 |
| Bone-Osteosarc | 41 | 117,874 | 0.2 | 0.48 | 0.51 | 8.9E-104 |
| Cervix-SCC | 18 | 84,315 | 0.14 | 0.35 | 0.36 | 7.2E-69 |
| Breast-LobularCa | 13 | 71,875 | 0.08 | 0.2 | 0.2 | 1.8E-19 |
| Kidney-ChRCC | 43 | 60,390 | 0.28 | 0.45 | 0.44 | 3.4E-51 |
| Thy-AdenoCA | 47 | 46,867 | 0.17 | 0.38 | 0.43 | 1.9E-54 |
| CNS-Oligo | 18 | 34,853 | 0.12 | 0.38 | 0.41 | 1.3E-66 |
| Lymph-NOS | 2 | 21,545 | 0.1 | 0.38 | 0.4 | 5.7E-162 |
| Myeloid-MPN | 24 | 18,753 | 0.2 | 0.36 | 0.39 | 1E-30 |
| Bone-Epith | 11 | 17,772 | 0.1 | 0.35 | 0.37 | 5E-84 |
| CNS-PiloAstro | 89 | 15,158 | 0.06 | 0.3 | 0.32 | 8.3E-66 |
| Myeloid-AML | 12 | 13,546 | 0.15 | 0.33 | 0.36 | 2.7E-36 |
| Cervix-AdenoCA | 2 | 5,998 | 0.11 | 0.24 | 0.24 | 2.3E-21 |
| Bone-Cart | 9 | 5,659 | 0.08 | 0.28 | 0.31 | 1E-45 |
| Breast-DCIS | 3 | 4,443 | 0.09 | 0.18 | 0.18 | 3.3E-15 |
| Myeloid-MDS | 1 | 403 | 0.03 | 0.05 | 0.04 | 0.09 |

**Supplementary Table 5**

**a)**
Different allele frequencies

| | 1kGP - chimpanzee | 1kGP - gorilla | chimpanzee - gorilla | 1kGP - tumor | chimpanzee - tumor | gorilla - tumor |
|---|---|---|---|---|---|---|
| **All segregating SNV** | 0.65 | 0.53 | 0.84 | 0.16 | 0.55 | 0.58 |
| **Singletons** | 0.49 | 0.25 | 0.53 | 0.15 | 0.47 | 0.43 |
| **Doubletons** | 0.38 | 0.19 | 0.46 | -0.02 | 0.45 | 0.39 |
| **Non-singletons** | 0.67 | 0.55 | 0.83 | 0.16 | 0.54 | 0.59 |
| **AF ≤ 0.05** | 0.63 | 0.57 | 0.71 | 0.2 | 0.53 | 0.44 |
| **AF > 0.05** | 0.65 | 0.58 | 0.79 | 0.2 | 0.5 | 0.59 |
| **AF ≤ 0.1** | 0.65 | 0.6 | 0.78 | 0.21 | 0.54 | 0.51 |
| **AF > 0.1** | 0.61 | 0.56 | 0.73 | 0.19 | 0.47 | 0.58 |

**b)**
Comparison between allele frequencies

| | Non-singletons vs Singletons | AF <= 5% vs AF > 5% | AF <= 10% vs AF > 10% |
|---|---|---|---|
| **1kGP** | 0.86 | 0.60 | 0.60 |
| **1kgp_69** | 0.70 | 0.71 | 0.73 |
| **1kgp_43** | 0.69 | 0.73 | 0.75 |
| **sgdp_50** | 0.72 | - | - |
| **chimpanzee** | 0.78 | 0.80 | 0.75 |
| **pp** | 0.35 | - | - |
| **ptv** | 0.23 | - | - |
| **pte** | 0.45 | - | - |
| **pts** | 0.61 | - | - |
| **ptt** | 0.75 | - | - |
| **gorilla** | 0.57 | 0.72 | 0.79 |
| **ggg** | 0.61 | - | - |
| **gbb** | -0.08 | - | - |
| **gbg** | 0.02 | - | - |

**c)**
Species-exclusive SNV

| | SNV exclusive of a species (not applied to tumor) |
|---|---|
| **1kGP exclusive vs Chimpanzee exclusive** | 0.55 |
| **1kGP exclusive vs Gorilla exclusive** | 0.43 |
| **1kGP exclusive vs tumor** | 0.16 |
| **Chimpanzee exclusive vs tumor** | 0.60 |
| **Gorilla exclusive vs tumor** | 0.62 |

**d)**
**Species-exclusive vs shared SNV**

| | shared |
|---|---|
| **1kGP exclusive vs shared 1kGP-chimp** | 0.64 |
| **Chimpanzee exclusive vs shared 1kGP-chimp** | 0.42 |
| **tumor vs shared 1kGP-chimp** | -0.02 |
| **1kGP exclusive vs shared 1kGP-gorilla** | 0.59 |
| **Gorilla exclusive vs shared 1kGP-gorilla** | 0.60 |
| **Tumor vs shared 1kGP-gorilla** | 0.13 |

**e)**
**Divergence**

| | Human-chimpanzee divergence | Human-Gorilla divergence |
|---|---|---|
| **1kGP** | 0.29 | 0.24 |
| **sgdp_50** | 0.26 | 0.22 |
| **Chimpanzee** | 0.20 | 0.19 |
| **chimp_50** | 0.20 | 0.20 |
| **Gorilla** | 0.24 | 0.13 |
| **Tumor** | 0.18 | 0.10 |

**Supplementary Table 6**

**a)**

SNV vs InDels

| | InDels 1kgp | InDels sgdp_50 | InDels Chimpanzee | InDels Gorilla | InDels Tumor |
|---|---|---|---|---|---|
| **SNV 1kgp** | 0.49 | 0.34 | 0.35 | 0.31 | -0.02 |
| **SNV sgdp_50** | 0.6 | 0.48 | 0.39 | 0.43 | 0.07 |
| **SNV Chimpanzee** | 0.43 | 0.26 | 0.49 | 0.53 | 0.39 |
| **SNV Gorilla** | 0.42 | 0.28 | 0.38 | 0.66 | 0.41 |
| **SNV Tumor** | 0.18 | 0.04 | 0.17 | 0.41 | 0.9 |

**b)**

InDels vs InDels

| | InDels 1kgp | InDels sgdp_50 | InDels Chimpanzee | InDels Gorilla | InDels Tumor |
|---|---|---|---|---|---|
| **InDels 1kgp** | - | - | - | - | - |
| **InDels sgdp_50** | 0.83 | - | - | - | - |
| **InDels Chimpanzee** | 0.69 | 0.64 | - | - | - |
| **InDels Gorilla** | 0.64 | 0.57 | 0.69 | - | - |
| **InDels Tumor** | 0.16 | 0.05 | 0.21 | 0.42 | - |

**Supplementary Table 7**

| Feature | Source | Type |
|---|---|---|
| GC-content | Calculated from the fraction of base pairs in a window passing our filters and being G or C in the human reference hg19 | Overlap |
| CpG | Calculated from the fraction of base pairs in a window passing our filters and being consecutive CpG in the human reference hg19 | Overlap |
| CpG islands | UCSC Genome Browser track: cpgIslandExt | Overlap |
| CpG methylation sites | UCSC Genome Browser track: CpG Methylation by Methyl 450K Bead Arrays from ENCODE/HAIB. Signal > 200 | Overlap |
| CpG hypermethylated in human/chimpanzee/gorilla | From Hernando-Herraez, I., Plos Genetics 2013. CpG sites marked as differently methylated in a species with no differences in other genera, and higher β-value in this species than in Orangutan | Overlap |
| CpG hypomethylated in human/chimpanzee/gorilla | From Hernando-Herraez, I., Plos Genetics 2013. CpG sites marked as differently methylated in a species with no differences in other genera, and lower β-value in this species than in Orangutan | Overlap |
| Genes | Protein-Coding Genes coordinates from ENSEMBL Genes 95 for human GRCh37.p13 | Overlap |
| Exons | Exon coordinates of Protein-Coding Genes from ENSEMBL Genes 95 for human GRCh37.p13 | Overlap |
| Recombination in Human (Male/Female/Average) | UCSC Genome Browser track: Recombination Rate from deCODE | Intensity |
| Recombination in Human PRDM9 DBS hotspots | From Pratto, F., Science 2014. Maps of Double Strand Breaks initiatied by PRDM9 binding sites. | Overlap |
| Recombination in Chimpanzee (Auton) | From Auton, A., Science 2012. Hotspots of recombination. Using *Pan troglodytes verus* | Overlap |
| Recombination in Chimpanzee (Stevison) | From Stevison, L., Mol Biol Evol 2015. Measures of rho/kb across the genome in Chimpanzee samples (*Pan troglodytes ellioti*) | Intensity |
| Recombination in Bonobo (Stevison) | From Stevison, L., Mol Biol Evol 2015. Measures of rho/kb across the genome in Bonobo samples | Intensity |
| Recombination in Gorilla (Stevison) | From Stevison, L., Mol Biol Evol 2015. Measures of rho/kb across the genome in Gorilla samples (*Gorilla gorilla gorilla*) | Intensity |
| Mappability of 24-mers | UCSC Genome Browser track: Mappability or Uniqueness of Reference Genome from ENCODE. | Intensity |
| Nucleosome Positioning | From Schones, D.E., Cell 2008. Downloaded from UCSC Genome Browser track: wgEncodeSydhNsomeGm12878Sig from ENCODE (Stanford, Snyder lab) using GM12878 cell line | Intensity |
| Methylation histone marks, H2AZ, CTCF and PolII binding | From Barski, A., Cell 2007. Measures of read-depth in CD4+ T cells | Intensity |
| Acetylation Histone Marks | From Wang, Z., Nature Genetics 2008. Measures of read-depth in CD4+ T cells | Intensity |
| Hi-C | From Lieberman-Aiden, E., Science 2009. | Intensity |
| Replication time | From Hansen, R.S., Proc. Natl Acad. Sci. 2010. Early-to-late ratio. | Intensity |
| Conservation | UCSC Genome Browser track: Conservation using 100 vertebrates - Vertebrate Multiz Alignment & Conservation (100 Species) - phyloP100wayAll | Intensity |
| Human ChromHMM chromatin states | From UCSC Genome Browser track: Broad ChromHMM - Chromatin State Segmentation by HMM from ENCODE/Broad in the GM12878 cell line. | Overlap |

**Supplementary Table 8**

**a)**

Fraction of CpG sites

| | | Fraction of ALL SNV | | | Fraction of Non-singleton SNV | | | Fraction of average heterozygous SNV per sample | |
|---|---|---|---|---|---|---|---|---|---|
| | **Count All** | **Non-CpG** | **CpG** | **Count All** | **Non-CpG** | **CpG** | **Count All** | **Non-CpG** | **CpG** |
| **1kGP** | 47,872,324 | 89.66% | 10.34% | 24,837,444 | 88.39% | 11.61% | - | - | - |
| **SGDP 50** | 10,651,708 | 89.04% | 10.96% | 5,845,807 | 89.06% | 10.94% | 912,555 | 89.28% | 10.72% |
| **Chimpanzee** | 26,872,437 | 89.91% | 10.09% | 18,741,493 | 89.69% | 10.31% | 1,408,912 | 89.67% | 10.33% |
| **Gorilla** | 9,813,559 | 89.22% | 10.78% | 8,431,675 | 89.25% | 10.75% | 1,289,016 | 89.59% | 10.41% |
| **Tumor** | 30,756,260 | 94.96% | 5.04% | - | - | - | - | - | - |

**b)**

Fraction of CpG doubletons

| | **% of shared between species doubletons at CpG sites** | **% of shared between species doubletons at CpG>T** |
|---|---|---|
| **sgdp_50 - African / non-African** | 6.91% | 7.10% |
| **Chimpanzee - pp / ptv / pte / pts /ptt** | 11.66% | 12.01% |
| **Chimpanzee - pp / ptv+pte / pts+ptt** | 7.62% | 7.89% |
| **Gorilla - ggg / gbb / gbg** | 6.10% | 6.13% |

**Supplementary Table 9**

**a)**

All SNV

| | Pearson's R | Poisson Regression R2 | Negative Binomial Regression R^2 |
|---|---|---|---|
| **1kGP** | 0.47 | 0.2 | 0.19 |
| **sgdp 50** | 0.38 | 0.14 | 0.13 |
| **Chimpanzee** | 0.23 | 0.06 | 0.06 |
| **Gorilla** | 0.18 | 0.03 | 0.03 |
| **Tumor** | 0.04 | 0 | 0 |

**b)**

non-CpG sites

| | Pearson's R | Poisson Regression R2 | Negative Binomial Regression R^2 |
|---|---|---|---|
| **1kGP** | 0.43 | 0.17 | 0.16 |
| **sgdp 50** | 0.35 | 0.12 | 0.11 |
| **Chimpanzee** | 0.26 | 0.07 | 0.07 |
| **Gorilla** | 0.21 | 0.05 | 0.05 |
| **Tumor** | 0.16 | 0.03 | 0.03 |

**c)**

CpG>T

| | Pearson's R | Poisson Regression R2 | Negative Binomial Regression R^2 |
|---|---|---|---|
| **1kGP** | 0.61 | 0.31 | 0.3 |
| **sgdp 50** | 0.59 | 0.29 | 0.27 |
| **Chimpanzee** | 0.6 | 0.3 | 0.28 |
| **Gorilla** | 0.56 | 0.27 | 0.25 |
| **Tumor** | -0.09 | 0.01 | 0.01 |

**Supplementary Table 10**

**a)**

Fraction of sites passing the 3-references filter

| | **non-CpG** | **CpG** |
|---|---|---|
| **1kGP** | 0.95 | 0.85 |
| **sgdp 50** | 0.94 | 0.84 |
| **Chimpanzee** | 0.86 | 0.81 |
| **Gorilla** | 0.7 | 0.66 |

**b)**

Correlation with mutation signatures

| | **SBS1** | **SBS5** | **10%*SBS1 + 90%SBS5** |
|---|---|---|---|
| **1kGP** | 0.4 | 0.92 | 0.97 |
| **sgdp 50** | 0.42 | 0.91 | 0.97 |
| **Chimpanzee** | 0.36 | 0.92 | 0.95 |
| **Gorilla** | 0.36 | 0.92 | 0.95 |
| **panTro5 sgdp 50** | 0.36 | 0.93 | 0.96 |
| **panTro5 Chimpanzee 50** | 0.36 | 0.92 | 0.95 |
| **Vervet Monkey** | 0.52 | 0.82 | 0.95 |
| **Mouse** | 0.13 | 0.82 | 0.74 |
| **Pig** | 0.4 | 0.75 | 0.83 |

**c)**

Correlation between mutation spectra

| | **1kGP** | **sgdp 50** | **Chimpanzee** | **Gorilla** | **panTro5 sgdp 50** | **panTro5 Chimpanzee 50** | **Vervet Monkey** | **Mouse** | **Pig** |
|---|---|---|---|---|---|---|---|---|---|
| **1kGP** | - | - | - | - | - | - | - | - | - |
| **sgdp 50** | 1 | - | - | - | - | - | - | - | - |
| **Chimpanzee** | 0.99 | 0.99 | - | - | - | - | - | - | - |
| **Gorilla** | 0.99 | 0.99 | 0.99 | - | - | - | - | - | - |
| **panTro5 sgdp 50** | 0.99 | 0.99 | 0.98 | 0.99 | - | - | - | - | - |
| **panTro5 Chimpanzee 50** | 0.98 | 0.98 | 0.99 | 0.98 | 0.99 | - | - | - | - |
| **Vervet Monkey** | 0.93 | 0.93 | 0.93 | 0.91 | 0.94 | 0.95 | - | - | - |
| **Mouse** | 0.83 | 0.82 | 0.82 | 0.79 | 0.77 | 0.78 | 0.87 | - | - |
| **Pig** | 0.82 | 0.82 | 0.82 | 0.8 | 0.82 | 0.83 | 0.85 | 0.8 | - |

**Supplementary Table 11**

| | ACC_A | ACT_A | CCG_A | GCG_A | TCG_A | ATT_A | TTA_A | ATG_G | CTG_G | GTG_G | TTG_G | proposed aetiology |
|---|---|---|---|---|---|---|---|---|---|---|---|---|
| **SBS7c** | | | | | | 0.051 | | | | | | UV-light damage |
| **SBS14** | | 0.083 | | | | | | | | | | Defective DNA mismatch repair (MSI) |
| **SBS22** | | | | | | | 0.072 | | | | | Aristolochic acid exposure |
| **SBS24** | | | | 0.066 | | | | | | | | Aflatoxin exposure |
| **SBS27** | | | | | | | 0.053 | | | | | Possible sequencing artefact |
| **SBS29** | 0.051 | | | | | | | | | | | Tobacco chewing habit |
| **SBS34** | | | | | | 0.316 | 0.272 | | | | | Unknown |
| **SBS38** | | | 0.051 | | | | | | | | | UV-light damage |
| **SBS41** | | | | | | 0.070 | 0.081 | | | | | Unknown |
| **SBS43** | | | | | | | | | | 0.241 | | Possible sequencing artefact |
| **SBS45** | | | 0.068 | | | | | | | | | Possible sequencing artefact |
| **SBS47** | | | | | | 0.068 | 0.156 | | | | | Possible sequencing artefact |
| **SBS48** | | | 0.137 | | 0.709 | | | | | | | Possible sequencing artefact |
| **SBS49** | | | 0.515 | | | | | | | | | Possible sequencing artefact |
| **SBS50** | 0.127 | 0.083 | | | | | | | | | | Possible sequencing artefact |
| **SBS51** | | | | | | | | | | 0.076 | | Possible sequencing artefact |
| **SBS52** | | | | | 0.134 | | | | | | | Possible sequencing artefact |
| **SBS53** | | | 0.068 | 0.051 | | | | | | | | Possible sequencing artefact |
| **SBS55** | | | | | | | | 0.145 | 0.113 | 0.299 | 0.080 | Possible sequencing artefact |
| **SBS60** | | | | | | | | | | 0.662 | | Possible sequencing artefact |
| **SBS85** | | | | | | 0.070 | 0.060 | | | | | Indirect effects of activation-induced cytidine deaminase (AID) induced somatic mutagenesis in lymphoid cells. |

**Supplementary Table 12**

**a)**
1kGP - chimpanzee

| | all | Skin-Melanoma | ColoRect-AdenoCA | Liver-HCC | Eso-AdenoCA | Lung-SCC | Stomach-AdenoCA | Panc-AdenoCA | Breast-AdenoCA | Lung-AdenoCA | Lymph-BNHL | Ovary-AdenoCA | Kidney-RCC | Head-SCC | Uterus-AdenoCA | Prost-AdenoCA | Bladder-TCC | CNS-GBM | Biliary-AdenoCA | Panc-Endocrine | Lymph-CLL | Bone-Leiomyo | CNS-Medullo | Bone-Osteosarc | Cervix-SCC | Breast-LobularCA | Kidney-ChRCC | Thy-AdenoCA | CNS-Oligo | Lymph-NOS | Myeloid-MPN | Bone-Epith | CNS-PiloAstro | Myeloid-AML | Bone-Cart | Cervix-AdenoCA | Breast-DCIS | Myeloid-MDS |
|---|---|---|---|---|---|---|---|---|---|---|---|---|---|---|---|---|---|---|---|---|---|---|---|---|---|---|---|---|---|---|---|---|---|---|---|---|---|---|
| **ALL** | 0.11 | 0.07 | 0.12 | 0.11 | 0.1 | 0.09 | 0.1 | 0.08 | 0.05 | 0.09 | 0.11 | 0.07 | 0.06 | 0.09 | 0.09 | 0.08 | 0.05 | - | 0.08 | 0.07 | 0.1 | 0.06 | 0.06 | 0.07 | 0.05 | - | - | - | - | 0.08 | - | 0.06 | - | - | - | - | - | - |
| **ACA>A** | - | - | - | - | - | - | - | - | - | - | - | - | - | - | - | - | - | - | - | - | - | - | - | - | - | - | - | - | - | - | - | - | - | - | - | - | - | - |
| **ACC>A** | - | - | - | - | - | - | - | - | - | - | - | - | - | - | - | - | - | - | - | - | - | - | - | - | - | - | - | - | - | - | - | - | - | - | - | - | - | - |
| **ACG>A** | - | - | - | - | - | - | - | - | - | - | - | - | - | - | - | - | - | - | - | - | - | - | - | - | - | - | - | - | - | - | - | - | - | - | - | - | - | - |
| **ACT>A** | - | - | - | - | - | - | - | - | - | - | - | - | - | - | - | - | - | - | - | - | - | - | - | - | - | - | - | - | - | - | - | - | - | - | - | - | - | - |
| **CCA>A** | - | - | - | - | - | - | - | - | - | - | - | - | - | - | - | - | - | - | - | - | - | - | - | - | - | - | - | - | - | - | - | - | - | - | - | - | - | - |
| **CCC>A** | - | - | - | - | - | - | - | - | - | - | - | - | - | - | - | - | - | - | - | - | - | - | - | - | - | - | - | - | - | - | - | - | - | - | - | - | - | - |
| **CCG>A** | - | - | - | - | - | - | - | - | - | - | - | - | - | - | - | - | - | - | - | - | - | - | - | - | - | - | - | - | - | - | - | - | - | - | - | - | - | - |
| **CCT>A** | - | - | - | - | - | - | - | - | - | - | - | - | - | - | - | - | - | - | - | - | - | - | - | - | - | - | - | - | - | - | - | - | - | - | - | - | - | - |
| **GCA>A** | - | - | - | - | - | - | - | - | - | - | - | - | - | - | - | - | - | - | - | - | - | - | - | - | - | - | - | - | - | - | - | - | - | - | - | - | - | - |
| **GCC>A** | - | - | - | - | - | - | - | - | - | - | - | - | - | - | - | - | - | - | - | - | - | - | - | - | - | - | - | - | - | - | - | - | - | - | - | - | - | - |
| **GCG>A** | - | - | - | - | - | - | - | - | - | - | - | - | - | - | - | - | - | - | - | - | - | - | - | - | - | - | - | - | - | - | - | - | - | - | - | - | - | - |
| **GCT>A** | - | - | - | - | - | - | - | - | - | - | - | - | - | - | - | - | - | - | - | - | - | - | - | - | - | - | - | - | - | - | - | - | - | - | - | - | - | - |
| **TCA>A** | - | - | - | - | - | - | - | - | - | - | - | - | - | - | - | - | - | - | - | - | - | - | - | - | - | - | - | - | - | - | - | - | - | - | - | - | - | - |
| **TCC>A** | - | - | - | - | - | - | - | - | - | - | - | - | - | - | - | - | - | - | - | - | - | - | - | - | - | - | - | - | - | - | - | - | - | - | - | - | - | - |
| **TCG>A** | - | - | - | - | - | - | - | - | - | - | - | - | - | - | - | - | - | - | - | - | - | - | - | - | - | - | - | - | - | - | - | - | - | - | - | - | - | - |
| **TCT>A** | - | - | - | - | - | - | - | - | - | - | - | - | - | - | - | - | - | - | - | - | - | - | - | - | - | - | - | - | - | - | - | - | - | - | - | - | - | - |
| **ACA>G** | 0.07 | - | - | 0.05 | - | 0.06 | - | - | - | - | - | - | - | - | - | - | - | - | - | - | - | - | - | - | - | - | - | - | - | - | - | - | - | - | - | - | - | - |
| **ACC>G** | - | - | - | - | - | - | - | - | - | - | - | - | - | - | - | - | - | - | - | - | - | - | - | - | - | - | - | - | - | - | - | - | - | - | - | - | - | - |
| **ACG>G** | - | - | - | - | - | - | - | - | - | - | - | - | - | - | - | - | - | - | - | - | - | - | - | - | - | - | - | - | - | - | - | - | - | - | - | - | - | - |
| **ACT>G** | 0.07 | - | - | - | - | - | - | - | - | - | - | - | - | - | - | - | - | - | - | - | - | - | - | - | - | - | - | - | - | - | - | - | - | - | - | - | - | - |
| **CCA>G** | 0.1 | - | - | 0.06 | 0.06 | 0.08 | - | - | - | 0.06 | 0.04 | - | - | 0.05 | - | - | - | - | - | - | - | - | - | - | - | - | - | - | - | - | - | - | - | - | - | - | - | - |
| **CCC>G** | 0.08 | - | - | - | - | 0.06 | - | - | - | - | - | - | - | - | - | - | - | - | - | - | - | - | - | - | - | - | - | - | - | - | - | - | - | - | - | - | - | - |
| **CCG>G** | - | - | - | - | - | - | - | - | - | - | - | - | - | - | - | - | - | - | - | - | - | - | - | - | - | - | - | - | - | - | - | - | - | - | - | - | - | - |
| **CCT>G** | 0.09 | 0.05 | - | 0.06 | - | 0.07 | - | - | - | 0.07 | - | - | - | - | - | - | - | - | - | - | - | - | - | - | - | - | - | - | - | - | - | - | - | - | - | - | - | - |
| **GCA>G** | 0.06 | - | - | - | - | - | - | - | - | - | - | - | - | - | - | - | - | - | - | - | - | - | - | - | - | - | - | - | - | - | - | - | - | - | - | - | - | - |
| **GCC>G** | - | - | - | - | - | - | - | - | - | - | - | - | - | - | - | - | - | - | - | - | - | - | - | - | - | - | - | - | - | - | - | - | - | - | - | - | - | - |
| **GCG>G** | - | - | - | - | - | - | - | - | - | - | - | - | - | - | - | - | - | - | - | - | - | - | - | - | - | - | - | - | - | - | - | - | - | - | - | - | - | - |
| **GCT>G** | 0.06 | - | - | - | - | - | - | - | - | - | - | - | - | - | - | - | - | - | - | - | - | - | - | - | - | - | - | - | - | - | - | - | - | - | - | - | - | - |
| **TCA>G** | 0.06 | - | - | - | - | - | - | - | - | - | - | - | - | - | - | - | - | - | - | - | - | - | - | - | - | - | - | - | - | - | - | - | - | - | - | - | - | - |
| **TCC>G** | 0.06 | - | - | - | - | - | - | - | - | - | - | - | - | - | - | - | - | - | - | - | - | - | - | - | - | - | - | - | - | - | - | - | - | - | - | - | - | - |
| **TCG>G** | - | - | - | - | - | - | - | - | - | - | - | - | - | - | - | - | - | - | - | - | - | - | - | - | - | - | - | - | - | - | - | - | - | - | - | - | - | - |
| **TCT>G** | 0.05 | - | 0.04 | - | 0.05 | - | - | - | - | - | - | - | - | - | - | - | - | - | - | - | - | - | - | - | - | - | - | - | - | - | - | - | - | - | - | - | - | - |
| **ACA>T** | 0.08 | - | 0.06 | 0.06 | 0.06 | 0.07 | - | 0.05 | - | 0.05 | 0.06 | - | - | 0.05 | - | - | - | - | - | - | - | - | - | - | - | - | - | - | - | - | - | - | - | - | - | - | - | - |
| **ACC>T** | - | - | - | - | - | - | - | - | - | - | - | - | - | - | - | - | - | - | - | - | - | - | - | - | - | - | - | - | - | - | - | - | - | - | - | - | - | - |
| **ACG>T** | 0.07 | - | 0.06 | - | - | - | - | - | - | - | - | - | - | - | - | - | - | - | - | - | - | - | - | - | - | - | - | - | - | - | - | - | - | - | - | - | - | - |
| **ACT>T** | - | - | - | - | - | - | - | - | - | - | - | - | - | - | - | - | - | - | - | - | - | - | - | - | - | - | - | - | - | - | - | - | - | - | - | - | - | - |
| **CCA>T** | - | - | - | - | - | - | - | - | - | - | - | - | - | - | - | - | - | - | - | - | - | - | - | - | - | - | - | - | - | - | - | - | - | - | - | - | - | - |
| **CCC>T** | - | - | - | - | - | - | - | - | - | - | - | - | - | - | - | - | - | - | - | - | - | - | - | - | - | - | - | - | - | - | - | - | - | - | - | - | - | - |
| **CCG>T** | - | - | - | - | - | - | - | - | - | - | - | - | - | - | - | - | - | - | - | - | - | - | - | - | - | - | - | - | - | - | - | - | - | - | - | - | - | - |
| **CCT>T** | - | - | - | - | - | - | - | - | - | - | - | - | - | - | - | - | - | - | - | - | - | - | - | - | - | - | - | - | - | - | - | - | - | - | - | - | - | - |
| **GCA>T** | - | - | - | - | - | - | - | - | - | - | - | - | - | - | - | - | - | - | - | - | - | - | - | - | - | - | - | - | - | - | - | - | - | - | - | - | - | - |
| **GCC>T** | - | - | - | - | - | - | - | - | - | - | - | - | - | - | - | - | - | - | - | - | - | - | - | - | - | - | - | - | - | - | - | - | - | - | - | - | - | - |
| **GCG>T** | - | - | - | - | - | - | - | - | - | - | - | - | - | - | - | - | - | - | - | - | - | - | - | - | - | - | - | - | - | - | - | - | - | - | - | - | - | - |
| **GCT>T** | - | - | - | - | - | - | - | - | - | - | - | - | - | - | - | - | - | - | - | - | - | - | - | - | - | - | - | - | - | - | - | - | - | - | - | - | - | - |
| **TCA>T** | - | - | - | - | - | - | - | - | - | - | - | - | - | - | - | - | - | - | - | - | - | - | - | - | - | - | - | - | - | - | - | - | - | - | - | - | - | - |
| **TCC>T** | 0.05 | 0.05 | 0.06 | - | - | - | - | - | - | - | - | - | - | - | - | - | - | - | - | - | - | - | - | - | - | - | - | - | - | - | - | - | - | - | - | - | - | - |
| **TCG>T** | 0.07 | 0.06 | 0.08 | - | - | - | - | - | - | - | - | - | - | - | - | - | - | - | - | - | - | - | - | - | - | - | - | - | - | - | - | - | - | - | - | - | - | - |
| **TCT>T** | - | - | - | - | - | - | - | - | - | - | - | - | - | - | - | - | - | - | - | - | - | - | - | - | - | - | - | - | - | - | - | - | - | - | - | - | - | - |
| **ATA>A** | - | - | - | - | - | - | - | - | - | - | - | - | - | - | - | - | - | - | - | - | - | - | - | - | - | - | - | - | - | - | - | - | - | - | - | - | - | - |
| **ATC>A** | - | - | - | - | - | - | - | - | - | - | - | - | - | - | - | - | - | - | - | - | - | - | - | - | - | - | - | - | - | - | - | - | - | - | - | - | - | - |
| **ATG>A** | - | - | - | - | - | - | - | - | - | - | - | - | - | - | - | - | - | - | - | - | - | - | - | - | - | - | - | - | - | - | - | - | - | - | - | - | - | - |
| **ATT>A** | - | - | - | - | - | - | - | - | - | - | - | - | - | - | - | - | - | - | - | - | - | - | - | - | - | - | - | - | - | - | - | - | - | - | - | - | - | - |
| **CTA>A** | - | - | - | - | - | - | - | - | - | - | - | - | - | - | - | - | - | - | - | - | - | - | - | - | - | - | - | - | - | - | - | - | - | - | - | - | - | - |
| **CTC>A** | - | - | - | - | - | - | - | - | - | - | - | - | - | - | - | - | - | - | - | - | - | - | - | - | - | - | - | - | - | - | - | - | - | - | - | - | - | - |
| **CTG>A** | - | - | - | - | - | - | - | - | - | - | - | - | - | - | - | - | - | - | - | - | - | - | - | - | - | - | - | - | - | - | - | - | - | - | - | - | - | - |
| **CTT>A** | - | - | - | - | - | - | - | - | - | - | - | - | - | - | - | - | - | - | - | - | - | - | - | - | - | - | - | - | - | - | - | - | - | - | - | - | - | - |
| **GTA>A** | - | - | - | - | - | - | - | - | - | - | - | - | - | - | - | - | - | - | - | - | - | - | - | - | - | - | - | - | - | - | - | - | - | - | - | - | - | - |
| **GTC>A** | - | - | - | - | - | - | - | - | - | - | - | - | - | - | - | - | - | - | - | - | - | - | - | - | - | - | - | - | - | - | - | - | - | - | - | - | - | - |
| **GTG>A** | - | - | - | - | - | - | - | - | - | - | - | - | - | - | - | - | - | - | - | - | - | - | - | - | - | - | - | - | - | - | - | - | - | - | - | - | - | - |
| **GTT>A** | - | - | - | - | - | - | - | - | - | - | - | - | - | - | - | - | - | - | - | - | - | - | - | - | - | - | - | - | - | - | - | - | - | - | - | - | - | - |
| **TTA>A** | - | - | - | - | - | - | - | - | - | - | - | - | - | - | - | - | - | - | - | - | - | - | - | - | - | - | - | - | - | - | - | - | - | - | - | - | - | - |
| **TTC>A** | - | - | - | - | - | - | - | - | - | - | - | - | - | - | - | - | - | - | - | - | - | - | - | - | - | - | - | - | - | - | - | - | - | - | - | - | - | - |
| **TTG>A** | - | - | - | - | - | - | - | - | - | - | - | - | - | - | - | - | - | - | - | - | - | - | - | - | - | - | - | - | - | - | - | - | - | - | - | - | - | - |
| **TTT>A** | - | - | - | - | - | - | - | - | - | - | - | - | - | - | - | - | - | - | - | - | - | - | - | - | - | - | - | - | - | - | - | - | - | - | - | - | - | - |
| **ATA>C** | 0.12 | 0.07 | 0.1 | 0.06 | 0.1 | 0.11 | 0.06 | 0.07 | - | 0.08 | 0.09 | 0.06 | - | 0.05 | - | - | 0.06 | - | - | - | - | - | - | - | - | - | - | - | - | - | - | - | - | - | - | - | - | - |
| **ATC>C** | 0.05 | - | - | - | - | - | - | - | - | - | - | - | - | - | - | - | - | - | - | - | - | - | - | - | - | - | - | - | - | - | - | - | - | - | - | - | - | - |
| **ATG>C** | 0.11 | - | 0.06 | 0.09 | 0.07 | 0.08 | - | - | - | 0.06 | 0.07 | - | - | - | - | - | - | - | - | - | - | - | - | - | - | - | - | - | - | - | - | - | - | - | - | - | - | - |
| **ATT>C** | 0.12 | 0.1 | 0.09 | 0.06 | 0.09 | 0.09 | 0.07 | 0.07 | - | 0.06 | 0.08 | 0.06 | - | 0.06 | - | - | - | 0.06 | - | - | - | - | - | - | - | - | - | - | - | - | - | - | - | - | - | - | - | - |
| **CTA>C** | 0.08 | 0.05 | - | 0.06 | 0.06 | 0.06 | 0.05 | - | - | - | 0.05 | - | - | - | - | - | - | - | - | - | - | - | - | - | - | - | - | - | - | - | - | - | - | - | - | - | - | - |
| **CTC>C** | - | - | - | - | - | - | - | - | - | - | - | - | - | - | - | - | - | - | - | - | - | - | - | - | - | - | - | - | - | - | - | - | - | - | - | - | - | - |
| **CTG>C** | 0.09 | - | - | 0.08 | 0.08 | 0.06 | 0.06 | - | - | - | 0.06 | - | - | - | - | - | - | - | - | - | - | - | - | - | - | - | - | - | - | - | - | - | - | - | - | - | - | - |
| **CTT>C** | 0.08 | 0.06 | 0.05 | 0.06 | 0.07 | - | 0.07 | - | - | - | 0.06 | - | - | - | - | - | - | - | - | - | - | - | - | - | - | - | - | - | - | - | - | - | - | - | - | - | - | - |
| **GTA>C** | - | - | - | - | - | - | - | - | - | - | - | - | - | - | - | - | - | - | - | - | - | - | - | - | - | - | - | - | - | - | - | - | - | - | - | - | - | - |
| **GTC>C** | - | - | - | - | - | - | - | - | - | - | - | - | - | - | - | - | - | - | - | - | - | - | - | - | - | - | - | - | - | - | - | - | - | - | - | - | - | - |
| **GTG>C** | 0.07 | - | - | 0.06 | - | - | - | - | - | - | - | - | - | - | - | - | - | - | - | - | - | - | - | - | - | - | - | - | - | - | - | - | - | - | - | - | - | - |
| **GTT>C** | 0.05 | - | - | - | - | - | - | - | - | - | - | - | - | - | - | - | - | - | - | - | - | - | - | - | - | - | - | - | - | - | - | - | - | - | - | - | - | - |
| **TTA>C** | 0.11 | 0.08 | 0.09 | 0.09 | 0.08 | 0.08 | 0.06 | 0.06 | - | 0.05 | 0.06 | - | - | 0.06 | - | - | - | - | - | - | - | - | - | - | - | - | - | - | - | - | - | - | - | - | - | - | - | - |
| **TTC>C** | - | - | - | - | - | - | - | - | - | - | - | - | - | - | - | - | - | - | - | - | - | - | - | - | - | - | - | - | - | - | - | - | - | - | - | - | - | - |
| **TTG>C** | 0.08 | - | 0.06 | 0.07 | 0.06 | 0.06 | - | - | - | - | - | - | - | - | - | - | - | - | - | - | - | - | - | - | - | - | - | - | - | - | - | - | - | - | - | - | - | - |
| **TTT>C** | 0.1 | 0.07 | 0.08 | 0.09 | 0.08 | 0.06 | 0.06 | - | - | - | 0.06 | - | - | - | - | - | - | - | - | - | - | - | - | - | - | - | - | - | - | - | - | - | - | - | - | - | - | - |
| **ATA>G** | - | - | - | - | - | - | - | - | - | - | - | - | - | - | - | - | - | - | - | - | - | - | - | - | - | - | - | - | - | - | - | - | - | - | - | - | - | - |
| **ATC>G** | - | - | - | - | - | - | - | - | - | - | - | - | - | - | - | - | - | - | - | - | - | - | - | - | - | - | - | - | - | - | - | - | - | - | - | - | - | - |
| **ATG>G** | - | - | - | - | - | - | - | - | - | - | - | - | - | - | - | - | - | - | - | - | - | - | - | - | - | - | - | - | - | - | - | - | - | - | - | - | - | - |
| **ATT>G** | - | - | - | - | - | - | - | - | - | - | - | - | - | - | - | - | - | - | - | - | - | - | - | - | - | - | - | - | - | - | - | - | - | - | - | - | - | - |
| **CTA>G** | - | - | - | - | - | - | - | - | - | - | - | - | - | - | - | - | - | - | - | - | - | - | - | - | - | - | - | - | - | - | - | - | - | - | - | - | - | - |
| **CTC>G** | - | - | - | - | - | - | - | - | - | - | - | - | - | - | - | - | - | - | - | - | - | - | - | - | - | - | - | - | - | - | - | - | - | - | - | - | - | - |
| **CTG>G** | - | - | - | - | - | - | - | - | - | - | - | - | - | - | - | - | - | - | - | - | - | - | - | - | - | - | - | - | - | - | - | - | - | - | - | - | - | - |
| **CTT>G** | - | - | - | - | - | - | - | - | - | - | - | - | - | - | - | - | - | - | - | - | - | - | - | - | - | - | - | - | - | - | - | - | - | - | - | - | - | - |
| **GTA>G** | - | - | - | - | - | - | - | - | - | - | - | - | - | - | - | - | - | - | - | - | - | - | - | - | - | - | - | - | - | - | - | - | - | - | - | - | - | - |
| **GTC>G** | - | - | - | - | - | - | - | - | - | - | - | - | - | - | - | - | - | - | - | - | - | - | - | - | - | - | - | - | - | - | - | - | - | - | - | - | - | - |
| **GTG>G** | - | - | - | - | - | - | - | - | - | - | - | - | - | - | - | - | - | - | - | - | - | - | - | - | - | - | - | - | - | - | - | - | - | - | - | - | - | - |
| **GTT>G** | - | - | - | - | - | - | - | - | - | - | - | - | - | - | - | - | - | - | - | - | - | - | - | - | - | - | - | - | - | - | - | - | - | - | - | - | - | - |
| **TTA>G** | - | - | - | - | - | - | - | - | - | - | - | - | - | - | - | - | - | - | - | - | - | - | - | - | - | - | - | - | - | - | - | - | - | - | - | - | - | - |
| **TTC>G** | 0.05 | - | - | - | - | - | - | - | - | - | - | - | - | - | - | - | - | - | - | - | - | - | - | - | - | - | - | - | - | - | - | - | - | - | - | - | - | - |
| **TTG>G** | 0.05 | - | - | - | - | - | - | - | - | - | - | - | - | - | - | - | - | - | - | - | - | - | - | - | - | - | - | - | - | - | - | - | - | - | - | - | - | - |
| **TTT>G** | - | - | - | - | - | - | - | - | - | - | - | - | - | - | - | - | - | - | - | - | - | - | - | - | - | - | - | - | - | - | - | - | - | - | - | - | - | - |

**b)**

1kGP - gorilla

| | all | Skin-Melanoma | ColoRect-AdenoCA | Liver-HCC | Eso-AdenoCA | Lung-SCC | Stomach-AdenoCA | Panc-AdenoCA | Breast-AdenoCA | Lung-AdenoCA | Lymph-BNHL | Ovary-AdenoCA | Kidney-RCC | Head-SCC | Uterus-AdenoCA | Prost-AdenoCA | Bladder-TCC | CNS-GBM | Biliary-AdenoCA | Panc-Endocrine | Lymph-CLL | Bone-Leiomyo | CNS-Medullo | Bone-Osteosarc | Cervix-SCC | Breast-LobularCA | Kidney-ChRCC | Thy-AdenoCA | CNS-Oligo | Lymph-NOS | Myeloid-MPN | Bone-Epith | CNS-PiloAstro | Myeloid-AML | Bone-Cart | Cervix-AdenoCA | Breast-DCIS | Myeloid-MDS |
|---|---|---|---|---|---|---|---|---|---|---|---|---|---|---|---|---|---|---|---|---|---|---|---|---|---|---|---|---|---|---|---|---|---|---|---|---|---|---|
| **ALL** | 0.11 | 0.08 | 0.12 | 0.1 | 0.09 | 0.09 | 0.09 | 0.07 | - | 0.09 | 0.1 | 0.06 | 0.06 | 0.09 | 0.09 | 0.07 | - | - | 0.07 | 0.06 | 0.1 | 0.07 | 0.06 | 0.07 | - | - | - | - | - | 0.08 | - | - | - | - | - | - | - | - |
| **ACA>A** | - | - | - | - | - | - | - | - | - | - | - | - | - | - | - | - | - | - | - | - | - | - | - | - | - | - | - | - | - | - | - | - | - | - | - | - | - | - |
| **ACC>A** | - | - | - | - | - | - | - | - | - | - | - | - | - | - | - | - | - | - | - | - | - | - | - | - | - | - | - | - | - | - | - | - | - | - | - | - | - | - |
| **ACG>A** | - | - | - | - | - | - | - | - | - | - | - | - | - | - | - | - | - | - | - | - | - | - | - | - | - | - | - | - | - | - | - | - | - | - | - | - | - | - |
| **ACT>A** | - | - | - | - | - | - | - | - | - | - | - | - | - | - | - | - | - | - | - | - | - | - | - | - | - | - | - | - | - | - | - | - | - | - | - | - | - | - |
| **CCA>A** | - | - | - | - | - | - | - | - | - | - | - | - | - | - | - | - | - | - | - | - | - | - | - | - | - | - | - | - | - | - | - | - | - | - | - | - | - | - |
| **CCC>A** | - | - | - | - | - | - | - | - | - | - | - | - | - | - | - | - | - | - | - | - | - | - | - | - | - | - | - | - | - | - | - | - | - | - | - | - | - | - |
| **CCG>A** | - | - | - | - | - | - | - | - | - | - | - | - | - | - | - | - | - | - | - | - | - | - | - | - | - | - | - | - | - | - | - | - | - | - | - | - | - | - |
| **CCT>A** | - | - | - | - | - | - | - | - | - | - | - | - | - | - | - | - | - | - | - | - | - | - | - | - | - | - | - | - | - | - | - | - | - | - | - | - | - | - |
| **GCA>A** | - | - | - | - | - | - | - | - | - | - | - | - | - | - | - | - | - | - | - | - | - | - | - | - | - | - | - | - | - | - | - | - | - | - | - | - | - | - |
| **GCC>A** | - | - | - | - | - | - | - | - | - | - | - | - | - | - | - | - | - | - | - | - | - | - | - | - | - | - | - | - | - | - | - | - | - | - | - | - | - | - |
| **GCG>A** | - | - | - | - | - | - | - | - | - | - | - | - | - | - | - | - | - | - | - | - | - | - | - | - | - | - | - | - | - | - | - | - | - | - | - | - | - | - |
| **GCT>A** | - | - | - | - | - | - | - | - | - | - | - | - | - | - | - | - | - | - | - | - | - | - | - | - | - | - | - | - | - | - | - | - | - | - | - | - | - | - |
| **TCA>A** | - | - | - | - | - | - | - | - | - | - | - | - | - | - | - | - | - | - | - | - | - | - | - | - | - | - | - | - | - | - | - | - | - | - | - | - | - | - |
| **TCC>A** | - | - | - | - | - | - | - | - | - | - | - | - | - | - | - | - | - | - | - | - | - | - | - | - | - | - | - | - | - | - | - | - | - | - | - | - | - | - |
| **TCG>A** | - | - | 0.01 | - | - | - | - | - | - | - | - | - | - | - | - | - | - | - | - | - | - | - | - | - | - | - | - | - | - | - | - | - | - | - | - | - | - | - |
| **TCT>A** | - | - | - | - | - | - | - | - | - | - | - | - | - | - | - | - | - | - | - | - | - | - | - | - | - | - | - | - | - | - | - | - | - | - | - | - | - | - |
| **ACA>G** | 0.07 | - | - | - | - | 0.06 | - | - | - | - | - | - | - | - | - | - | - | - | - | - | - | - | - | - | - | - | - | - | - | - | - | - | - | - | - | - | - | - |
| **ACC>G** | - | - | - | - | - | - | - | - | - | - | - | - | - | - | - | - | - | - | - | - | - | - | - | - | - | - | - | - | - | - | - | - | - | - | - | - | - | - |
| **ACG>G** | - | - | - | - | - | 0.01 | - | - | - | - | - | - | - | - | - | - | - | - | - | - | - | - | - | - | - | - | - | - | - | - | - | - | - | - | - | - | - | - |
| **ACT>G** | 0.06 | - | - | - | - | - | - | - | - | - | - | - | - | - | - | - | - | - | - | - | - | - | - | - | - | - | - | - | - | - | - | - | - | - | - | - | - | - |
| **CCA>G** | 0.08 | - | - | 0.06 | - | 0.06 | - | - | - | - | - | - | - | - | - | - | - | - | - | - | - | - | - | - | - | - | - | - | - | - | - | - | - | - | - | - | - | - |
| **CCC>G** | 0.08 | - | - | - | - | 0.06 | - | - | - | 0.05 | - | - | - | - | - | - | - | - | - | - | - | - | - | - | - | - | - | - | - | - | - | - | - | - | - | - | - | - |
| **CCG>G** | - | 0 | - | - | - | - | - | - | - | - | - | - | - | - | - | - | - | - | - | - | - | - | - | - | - | - | - | - | - | - | - | - | - | - | - | - | - | - |
| **CCT>G** | 0.1 | 0.06 | - | 0.06 | - | 0.08 | - | - | - | 0.08 | - | - | - | - | - | - | - | - | - | - | - | - | - | - | - | - | - | - | - | - | - | - | - | - | - | - | - | - |
| **GCA>G** | 0.06 | - | - | - | - | - | - | - | - | - | - | - | - | - | - | - | - | - | - | - | - | - | - | - | - | - | - | - | - | - | - | - | - | - | - | - | - | - |
| **GCC>G** | - | - | - | - | - | - | - | - | - | - | - | - | - | - | - | - | - | - | - | - | - | - | - | - | - | - | - | - | - | - | - | - | - | - | - | - | - | - |
| **GCG>G** | - | 0.02 | - | 0.01 | 0.02 | 0.01 | 0.03 | 0.03 | 0.02 | 0.01 | - | 0.01 | 0.03 | 0.02 | - | - | 0.03 | - | - | 0.03 | 0.03 | - | - | 0.03 | - | 0.03 | - | - | 0.03 | 0.03 | - | - | - | - | - | - | 0.03 | 0.03 |
| **GCT>G** | 0.05 | - | - | - | - | - | - | - | - | - | - | - | - | - | - | - | - | - | - | - | - | - | - | - | - | - | - | - | - | - | - | - | - | - | - | - | - | - |
| **TCA>G** | 0.06 | - | - | 0.07 | - | - | - | - | - | - | - | - | - | - | - | - | - | - | - | - | - | - | - | - | - | - | - | - | - | - | - | - | - | - | - | - | - | - |
| **TCC>G** | 0.08 | - | - | - | - | 0.06 | - | - | - | - | - | - | - | - | - | - | - | - | - | - | - | - | - | - | - | - | - | - | - | - | - | - | - | - | - | - | - | - |
| **TCG>G** | - | - | - | - | - | 0.02 | - | - | - | - | - | - | - | - | - | - | - | - | - | - | - | - | - | - | - | - | - | - | - | - | - | - | - | - | - | - | - | - |
| **TCT>G** | 0.08 | 0.07 | 0.07 | 0.07 | 0.07 | 0.06 | - | 0.06 | - | - | 0.06 | - | - | - | - | - | - | - | - | - | - | - | - | - | - | - | - | - | - | - | - | - | - | - | - | - | - | - |
| **ACA>T** | 0.08 | - | 0.06 | 0.06 | 0.06 | 0.06 | - | 0.05 | - | 0.05 | 0.06 | - | - | 0.05 | - | - | - | - | - | - | - | - | - | - | - | - | - | - | - | - | - | - | - | - | - | - | - | - |
| **ACC>T** | - | - | - | - | - | - | - | - | - | - | - | - | - | - | - | - | - | - | - | - | - | - | - | - | - | - | - | - | - | - | - | - | - | - | - | - | - | - |
| **ACG>T** | - | - | - | - | - | - | - | - | - | - | - | - | - | - | - | - | - | - | - | - | - | - | - | - | - | - | - | - | - | - | - | - | - | - | - | - | - | - |
| **ACT>T** | - | - | - | - | - | - | - | - | - | - | - | - | - | - | - | - | - | - | - | - | - | - | - | - | - | - | - | - | - | - | - | - | - | - | - | - | - | - |
| **CCA>T** | - | - | - | - | - | - | - | - | - | - | - | - | - | - | - | - | - | - | - | - | - | - | - | - | - | - | - | - | - | - | - | - | - | - | - | - | - | - |
| **CCC>T** | - | - | - | - | - | - | - | - | - | - | - | - | - | - | - | - | - | - | - | - | - | - | - | - | - | - | - | - | - | - | - | - | - | - | - | - | - | - |
| **CCG>T** | - | - | - | - | - | - | - | - | - | - | - | - | - | - | - | - | - | - | - | - | - | - | - | - | - | - | - | - | - | - | - | - | - | - | - | - | - | - |
| **CCT>T** | - | - | - | - | - | - | - | - | - | - | - | - | - | - | - | - | - | - | - | - | - | - | - | - | - | - | - | - | - | - | - | - | - | - | - | - | - | - |
| **GCA>T** | - | - | - | - | - | - | - | - | - | - | - | - | - | - | - | - | - | - | - | - | - | - | - | - | - | - | - | - | - | - | - | - | - | - | - | - | - | - |
| **GCC>T** | - | - | - | - | - | - | - | - | - | - | - | - | - | - | - | - | - | - | - | - | - | - | - | - | - | - | - | - | - | - | - | - | - | - | - | - | - | - |
| **GCG>T** | - | - | - | - | - | - | - | - | - | - | - | - | - | - | - | - | - | - | - | - | - | - | - | - | - | - | - | - | - | - | - | - | - | - | - | - | - | - |
| **GCT>T** | - | - | - | - | - | - | - | - | - | - | - | - | - | - | - | - | - | - | - | - | - | - | - | - | - | - | - | - | - | - | - | - | - | - | - | - | - | - |
| **TCA>T** | 0.08 | 0.08 | 0.07 | 0.07 | 0.06 | 0.05 | - | - | - | - | 0.07 | - | - | - | - | - | - | - | - | - | - | - | - | - | - | - | - | - | - | - | - | - | - | - | - | - | - | - |
| **TCC>T** | 0.05 | - | 0.06 | - | - | - | - | - | - | - | - | - | - | - | - | - | - | - | - | - | - | - | - | - | - | - | - | - | - | - | - | - | - | - | - | - | - | - |
| **TCG>T** | - | - | - | - | - | - | - | - | - | - | - | - | - | - | - | - | - | - | - | - | - | - | - | - | - | - | - | - | - | - | - | - | - | - | - | - | - | - |
| **TCT>T** | - | - | 0.05 | - | - | - | - | - | - | - | - | - | - | - | - | - | - | - | - | - | - | - | - | - | - | - | - | - | - | - | - | - | - | - | - | - | - | - |
| **ATA>A** | - | - | - | - | - | - | - | - | - | - | - | - | - | - | - | - | - | - | - | - | - | - | - | - | - | - | - | - | - | - | - | - | - | - | - | - | - | - |
| **ATC>A** | - | - | - | - | - | - | - | - | - | - | - | - | - | - | - | - | - | - | - | - | - | - | - | - | - | - | - | - | - | - | - | - | - | - | - | - | - | - |
| **ATG>A** | - | - | - | - | - | - | - | - | - | - | - | - | - | - | - | - | - | - | - | - | - | - | - | - | - | - | - | - | - | - | - | - | - | - | - | - | - | - |
| **ATT>A** | - | - | - | - | - | - | - | - | - | - | - | - | - | - | - | - | - | - | - | - | - | - | - | - | - | - | - | - | - | - | - | - | - | - | - | - | - | - |
| **CTA>A** | - | - | - | - | - | - | - | - | - | - | - | - | - | - | - | - | - | - | - | - | - | - | - | - | - | - | - | - | - | - | - | - | - | - | - | - | - | - |
| **CTC>A** | - | - | - | - | - | - | - | - | - | - | - | - | - | - | - | - | - | - | - | - | - | - | - | - | - | - | - | - | - | - | - | - | - | - | - | - | - | - |
| **CTG>A** | - | - | - | - | - | - | - | - | - | - | - | - | - | - | - | - | - | - | - | - | - | - | - | - | - | - | - | - | - | - | - | - | - | - | - | - | - | - |
| **CTT>A** | - | - | - | - | - | - | - | - | - | - | - | - | - | - | - | - | - | - | - | - | - | - | - | - | - | - | - | - | - | - | - | - | - | - | - | - | - | - |
| **GTA>A** | - | - | - | - | - | - | - | - | - | - | - | - | - | - | - | - | - | - | - | - | - | - | - | - | - | - | - | - | - | - | - | - | - | - | - | - | - | - |
| **GTC>A** | - | - | - | - | - | - | - | - | - | - | - | - | - | - | - | - | - | - | - | - | - | - | - | - | - | - | - | - | - | - | - | - | - | - | - | - | - | - |
| **GTG>A** | - | - | - | - | - | - | - | - | - | - | - | - | - | - | - | - | - | - | - | - | - | - | - | - | - | - | - | - | - | - | - | - | - | - | - | - | - | - |
| **GTT>A** | - | - | - | - | - | - | - | - | - | - | - | - | - | - | - | - | - | - | - | - | - | - | - | - | - | - | - | - | - | - | - | - | - | - | - | - | - | - |
| **TTA>A** | - | - | - | - | - | - | - | - | - | - | - | - | - | - | - | - | - | - | - | - | - | - | - | - | - | - | - | - | - | - | - | - | - | - | - | - | - | - |
| **TTC>A** | - | - | - | - | - | - | - | - | - | - | - | - | - | - | - | - | - | - | - | - | - | - | - | - | - | - | - | - | - | - | - | - | - | - | - | - | - | - |
| **TTG>A** | - | - | - | - | - | - | - | - | - | - | - | - | - | - | - | - | - | - | - | - | - | - | - | - | - | - | - | - | - | - | - | - | - | - | - | - | - | - |
| **TTT>A** | - | - | - | - | - | - | - | - | - | - | - | - | - | - | - | - | - | - | - | - | - | - | - | - | - | - | - | - | - | - | - | - | - | - | - | - | - | - |
| **ATA>C** | 0.14 | 0.08 | 0.12 | 0.06 | 0.11 | 0.13 | 0.07 | 0.09 | - | 0.09 | 0.1 | 0.07 | - | - | - | - | 0.07 | - | - | - | - | - | - | - | - | - | - | - | - | - | - | - | - | - | - | - | - | - |
| **ATC>C** | 0.05 | - | - | - | - | - | - | - | - | - | - | - | - | - | - | - | - | - | - | - | - | - | - | - | - | - | - | - | - | - | - | - | - | - | - | - | - | - |
| **ATG>C** | 0.13 | 0.07 | 0.07 | 0.11 | 0.09 | 0.1 | - | 0.06 | - | 0.06 | 0.08 | - | - | 0.06 | - | - | - | - | - | - | - | - | - | - | - | - | - | - | - | - | - | - | - | - | - | - | - | - |
| **ATT>C** | 0.14 | 0.12 | 0.11 | 0.07 | 0.11 | 0.11 | 0.08 | 0.08 | - | 0.07 | 0.1 | 0.07 | - | 0.06 | - | - | - | 0.07 | - | - | 0.06 | - | - | - | - | - | - | - | - | - | - | - | - | - | - | - | - | - |
| **CTA>C** | 0.08 | - | - | 0.07 | 0.06 | 0.06 | - | - | - | - | - | - | - | - | - | - | - | - | - | - | - | - | - | - | - | - | - | - | - | - | - | - | - | - | - | - | - | - |
| **CTC>C** | - | - | - | - | - | - | - | - | - | - | - | - | - | - | - | - | - | - | - | - | - | - | - | - | - | - | - | - | - | - | - | - | - | - | - | - | - | - |
| **CTG>C** | 0.1 | - | - | 0.09 | 0.09 | 0.07 | 0.07 | - | - | 0.06 | 0.07 | - | - | - | - | - | - | - | - | - | - | - | - | - | - | - | - | - | - | - | - | - | - | - | - | - | - | - |
| **CTT>C** | 0.09 | 0.08 | 0.06 | 0.07 | 0.08 | 0.06 | 0.08 | 0.05 | - | - | 0.07 | - | - | - | - | - | - | - | - | - | - | - | - | - | - | - | - | - | - | - | - | - | - | - | - | - | - | - |
| **GTA>C** | - | - | - | - | - | - | - | - | - | - | - | - | - | - | - | - | - | - | - | - | - | - | - | - | - | - | - | - | - | - | - | - | - | - | - | - | - | - |
| **GTC>C** | - | - | - | - | - | - | - | - | - | - | - | - | - | - | - | - | - | - | - | - | - | - | - | - | - | - | - | - | - | - | - | - | - | - | - | - | - | - |
| **GTG>C** | 0.07 | - | - | 0.06 | 0.05 | - | - | - | - | - | - | - | - | - | - | - | - | - | - | - | - | - | - | - | - | - | - | - | - | - | - | - | - | - | - | - | - | - |
| **GTT>C** | 0.05 | - | - | - | - | - | - | - | - | - | - | - | - | - | - | - | - | - | - | - | - | - | - | - | - | - | - | - | - | - | - | - | - | - | - | - | - | - |
| **TTA>C** | 0.11 | 0.08 | 0.09 | 0.09 | 0.08 | 0.08 | 0.05 | 0.06 | - | - | 0.06 | - | - | 0.06 | - | - | - | - | - | - | - | - | - | - | - | - | - | - | - | - | - | - | - | - | - | - | - | - |
| **TTC>C** | - | - | - | - | - | - | - | - | - | - | - | - | - | - | - | - | - | - | - | - | - | - | - | - | - | - | - | - | - | - | - | - | - | - | - | - | - | - |
| **TTG>C** | 0.09 | 0.05 | 0.06 | 0.07 | 0.07 | 0.07 | - | - | - | - | - | - | - | - | - | - | - | - | - | - | - | - | - | - | - | - | - | - | - | - | - | - | - | - | - | - | - | - |
| **TTT>C** | 0.12 | 0.1 | 0.1 | 0.11 | 0.11 | 0.08 | 0.08 | 0.06 | - | - | 0.08 | - | - | 0.06 | - | - | - | - | - | - | - | - | - | - | - | - | - | - | - | - | - | - | - | - | - | - | - | - |
| **ATA>G** | - | - | - | - | - | - | - | - | - | - | - | - | - | - | - | - | - | - | - | - | - | - | - | - | - | - | - | - | - | - | - | - | - | - | - | - | - | - |
| **ATC>G** | - | - | - | - | - | - | - | - | - | - | - | - | - | - | - | - | - | - | - | - | - | - | - | - | - | - | - | - | - | - | - | - | - | - | - | - | - | - |
| **ATG>G** | - | - | - | - | - | - | - | - | - | - | - | - | - | - | - | - | - | - | - | - | - | - | - | - | - | - | - | - | - | - | - | - | - | - | - | - | - | - |
| **ATT>G** | - | - | - | - | - | - | - | - | - | - | - | - | - | - | - | - | - | - | - | - | - | - | - | - | - | - | - | - | - | - | - | - | - | - | - | - | - | - |
| **CTA>G** | - | - | - | - | - | - | - | - | - | - | - | - | - | - | - | - | - | - | - | - | - | - | - | - | - | - | - | - | - | - | - | - | - | - | - | - | - | - |
| **CTC>G** | - | - | - | - | - | - | - | - | - | - | - | - | - | - | - | - | - | - | - | - | - | - | - | - | - | - | - | - | - | - | - | - | - | - | - | - | - | - |
| **CTG>G** | - | - | - | - | - | - | - | - | - | - | - | - | - | - | - | - | - | - | - | - | - | - | - | - | - | - | - | - | - | - | - | - | - | - | - | - | - | - |
| **CTT>G** | - | - | - | - | - | - | - | - | - | - | - | - | - | - | - | - | - | - | - | - | - | - | - | - | - | - | - | - | - | - | - | - | - | - | - | - | - | - |
| **GTA>G** | - | - | - | - | - | - | - | - | - | - | - | - | - | - | - | - | - | - | - | - | - | - | - | - | - | - | - | - | - | - | - | - | - | - | - | - | - | - |
| **GTC>G** | - | - | - | - | - | - | - | - | - | - | - | - | - | - | - | - | - | - | - | - | - | - | - | - | - | - | - | - | - | - | - | - | - | - | - | - | - | - |
| **GTG>G** | - | - | - | - | - | - | - | - | - | - | - | - | - | - | - | - | - | - | - | - | - | - | - | - | - | - | - | - | - | - | - | - | - | - | - | - | - | - |
| **GTT>G** | - | - | - | - | - | - | - | - | - | - | - | - | - | - | - | - | - | - | - | - | - | - | - | - | - | - | - | - | - | - | - | - | - | - | - | - | - | - |
| **TTA>G** | - | - | - | - | - | - | - | - | - | - | - | - | - | - | - | - | - | - | - | - | - | - | - | - | - | - | - | - | - | - | - | - | - | - | - | - | - | - |
| **TTC>G** | 0.07 | 0.04 | - | - | 0.05 | - | - | - | - | - | - | - | - | - | - | - | - | - | - | - | - | - | - | - | - | - | - | - | - | - | - | - | - | - | - | - | - | - |
| **TTG>G** | - | - | - | - | - | - | - | - | - | - | - | - | - | - | - | - | - | - | - | - | - | - | - | - | - | - | - | - | - | - | - | - | - | - | - | - | - | - |
| **TTT>G** | - | - | - | - | - | - | - | - | - | - | - | - | - | - | - | - | - | - | - | - | - | - | - | - | - | - | - | - | - | - | - | - | - | - | - | - | - | - |
